# Ribosome profiling reveals widespread translational control across developmental stages in the brown alga *Ectocarpus*

**DOI:** 10.64898/2026.04.17.719139

**Authors:** Jia Xuan Leong, Josué Barrera-Redondo, Fabian Haas, Cátia Igreja, Susana M Coelho

## Abstract

Translation represents a central regulatory layer of gene expression, enabling rapid and dynamic modulation of protein output. While its importance in development is well established, it remains poorly characterized outside animals and land plants. Brown algae, and in particular *Ectocarpus*, offer a compelling system to investigate translational regulation in development, having evolved multicellularity independently of other major eukaryotic lineages and exhibiting a complex life cycle with distinct haploid and diploid developmental programs. Although transcriptional regulation contributes to *Ectocarpus* development, the striking similarity of transcriptomes across life cycle stages indicates that transcription alone is insufficient to explain their distinct developmental identities, pointing to a major role for post-transcriptional regulation. Here, we establish ribosome profiling in *Ectocarpus* and define its translatome across key developmental stages. We uncover widespread stage- and sex-specific translational regulation, with post-transcriptional buffering emerging as the predominant mode of control. Genes subject to translational regulation exhibit optimized codon usage, broader transcriptional profiles and shorter 3′ UTRs. These genomic features are associated with variation in translation efficiency. Collectively, these results provide the first comprehensive characterization of the *Ectocarpus* translatome and highlight a major role for translational regulation in developmental gene expression in an independently evolved multicellular lineage.

## Introduction

Translation is a conserved biological process that synthesizes proteins by decoding messenger RNA (mRNA) on the ribosome. This process involves transfer RNAs (tRNAs) and numerous auxiliary factors and proceeds through three mechanistically distinct stages - initiation, elongation, and termination - each subject to diverse and context-dependent regulation (1–6).

While gene expression is often inferred from mRNA levels, translational regulation can generate substantial differences between mRNA abundance and protein output for individual genes (7, 8). Recent advances in measuring translation independently of transcript abundance have established translational regulation as a central mechanism that shapes the proteome (9, 10).

Among the diverse outcomes of translational regulation, post-transcriptional buffering has emerged as an important mechanism that maintains protein homeostasis despite variation in mRNA levels (8, 11–13). This process involves adjustments in ribosome occupancy that offset transcriptional variation, so that changes in mRNA abundance do not translate into proportional differences in protein output. Transcripts subject to translational buffering can be characterized by specific structural features in their 5’ untranslated regions (UTRs) or exhibit preferential association with a subset of tRNAs during translation, as shown in studies of ERα depletion in human cancer cells (12).

Translational regulation plays essential roles in development and cellular differentiation in animals and land plants (14–17). For example, the oocyte-to-embryo transition in *Drosophila* involves extensive remodeling of the translatome in the absence of transcription and mRNA degradation (18). In *Arabidopsis*, global translational regulation occurs during the early phase of seed maturation under the control of the key regulator ABSCISIC ACID INSENSITIVE3 (ABI3) (19). Whether comparable roles extend to other multicellular and evolutionarily distant eukaryotes, however, remains unclear.

Brown algae represent a deeply divergent and largely unexplored eukaryotic lineage, that provides a unique opportunity to examine the evolution and developmental roles of gene regulatory mechanisms. They are distantly related to animals, plants, and fungi, and have independently evolved complex multicellularity (20–22). Among them, *Ectocarpus* sp.7 (hereafter *Ectocarpus*) has emerged as a tractable model for evolutionary and developmental studies (22, 23). Notably, the *Ectocarpus* life cycle alternates between distinct multicellular sporophyte and gametophyte generations (24, 25), requiring precise gene regulation to deploy the appropriate developmental program at each stage (22). Transcriptome analyses have revealed a high degree of similarity in mRNA profiles across *Ectocarpus* developmental stages and tissues, indicating that transcriptional regulation alone cannot account for developmental and functional diversity (26–28). These observations point to a potential role for post-transcriptional mechanisms, including translational regulation, in developmental programs across alternation of generations. However, the translational landscape of *Ectocarpus* remains uncharacterized, and the contribution of translational regulation to its life cycle is unknown.

Here, we combine ribosome profiling (Ribo-seq) with transcriptomic (RNA-seq) and proteomic analyses to investigate translational regulation in *Ectocarpus* across developmental stages and sexes. Integrating these complementary datasets enables direct comparison of transcriptional and translational outputs to identify genes subject to translational control and buffering. This work establishes the first ribosome profiling framework in brown algae and provides a comprehensive view of translational regulation across the *Ectocarpus* life cycle, highlighting brown algae as an evolutionarily distinct system that extends gene-regulatory models beyond animals and land plants.

## Results

### Tissue-specific ionic conditions enable high-quality ribosome profiling in *Ectocarpus*

To assess the contribution of translation to gene-expression regulation across *Ectocarpus* developmental stages, we established a high-quality Ribo-seq protocol using tissue from three life-cycle stages: partheno-sporophytes (pSP) and female and male gametophytes (GA) (**Fig. 1A, Table S1**). These samples enabled analysis of sex-specific control of protein synthesis (female vs. male GA) and comparison of translational regulation between life-cycle stages (GA vs. pSP).

**Figure 1.**
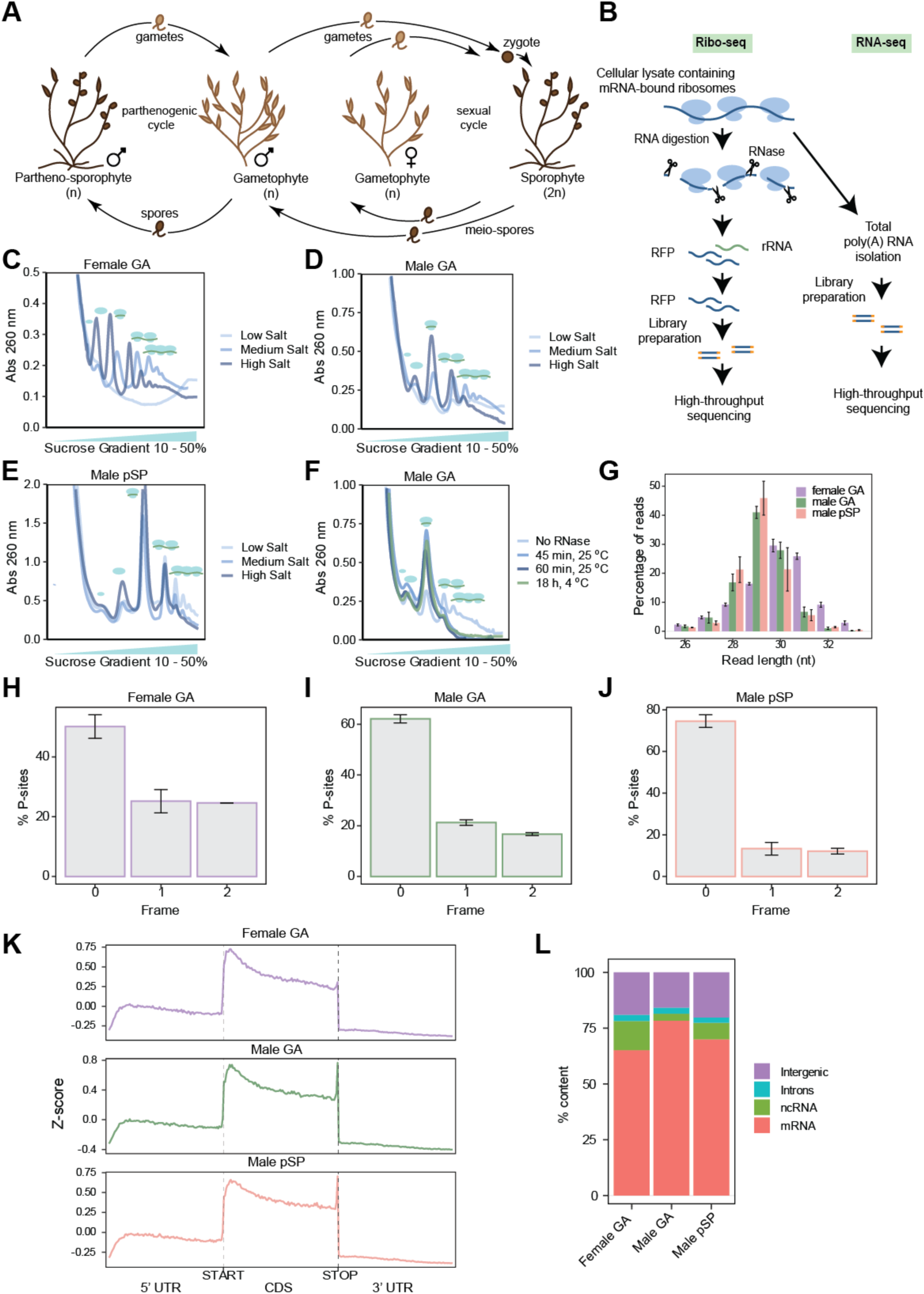
Establishment of ribosome profiling in *Ectocarpus*. **(A)** Life cycle of *Ectocarpus*. During the sexual cycle, haploid male and female gametophytes (GA) release gametes, which undergo syngamy to form a zygote which develops into a diploid sporophyte (SP). Upon maturity, sporophytes release meio-spores which develop into GA. Male and female gametes, when unfertilized, can undergo parthenogenesis and develop into haploid partheno-sporophytes (pSP). **(B)** Overview of the ribosome profiling workflow including parallel Ribo-seq and RNA-seq library preparation. RFP, ribosome footprints. **(C–E)** Polysome profiles (UV absorbance at 260 nm) from sucrose gradient fractionation of **(C)** female GA, **(D)** male GA, and **(E)** pSP lysates under low (50 mM NaCl, 5 mM MgCl₂), medium (200 mM NaCl, 20 mM MgCl₂), and high (700 mM NaCl, 70 mM MgCl₂) salt conditions. Peaks from left to right correspond to the small (40S) and large (60S) ribosomal subunits, 80S monosomes, disomes, and polysomes, as indicated by the schematic icons above each peak. **(F)** Polysome collapse assay with different RNase I digestion conditions (no RNase control, 1 U μL^-1^ RNase I 45 min and 60 min at 25°C, and 18 h at 4°C) for male GA lysate. **(G)** RFP length distribution across female GA, male GA and male pSP datasets after quality filtering of the Ribo-seq libraries prepared from lysates digested with 4.25 U μL^-1^ of RNase I. Predominant footprint sizes range from 28–31 nts. **(H–J)** Reading frame distribution scores of P-sites for **(H)** female GA, **(I)** male GA, and **(J)** male pSP, showing strong frame 0 bias. **(K)** Metagene profiles of ribosome footprint density across scaled transcript regions (5′ UTR, CDS, 3′ UTR) for each developmental stage. Ribosome footprint density was normalized per gene by computing Z-scores (score − mean) / standard deviation within each transcript and condition, then averaged across all transcripts to generate the metagene profile. **(L)** Distribution of mapped Ribo-seq reads across genomic features for each developmental stage.

Isolation of ribosome-protected RNA fragments depends on lysis conditions, nucleolytic digestion, ribosome purification, and precise selection of ribosome footprints (RFPs) for library preparation (**Fig. 1B**) (29, 30). To establish a robust Ribo-seq protocol for brown algae, we first optimized lysis conditions for efficient recovery of ribosomal complexes from different *Ectocarpus* tissues. Because ribosome integrity is sensitive to ionic composition, we varied Mg^2+^ and Na^+^ concentrations in the lysis buffer. Polysome profiling revealed tissue-specific requirements for salt conditions (**Fig. 1C–E; Supplementary Fig. 1A–C**).

In GA, only medium salt conditions (200 mM NaCl, 20 mM MgCl₂) preserved intact monosomes and polysomes (**Fig. 1C-D**), whereas pSP yielded stable ribosomal complexes under both low salt (50 mM NaCl, 5 mM MgCl₂) and medium salt conditions (**Fig. 1E**). High salt buffers (700 mM NaCl, 70 mM MgCl2) disrupted ribosomal complexes integrity in all tissues, as indicated by increased free 40S/60S ribosomal subunits, reduced polysomes, and 18S rRNA degradation (**Fig. 1C–E; Supplementary Fig. 1A–C)**.

Independent variation of NaCl and MgCl₂ concentrations in GA samples showed that 50 mM NaCl impaired ribosome recovery regardless of Mg²⁺ concentration, whereas 200 mM NaCl preserved complex integrity (**Supplementary Fig. 1D-E**). Increasing Mg²⁺ further improved ribosome stability, although less strongly than Na⁺. These results indicate distinct ionic requirements for ribosome isolation in GA and pSP tissues, potentially reflecting differences in intracellular ion composition or lysis efficiency.

Based on the influence of the ionic strength of the lysis buffer in the isolation of ribosomal complexes with high quality, all subsequent experiments were conducted under low and medium salt conditions for the pSP and GA tissues, respectively.

We next subjected algal lysates to nucleolytic digestion, a critical step in ribosome profiling that enriches for RFPs. We used RNase I, a broadly employed enzyme lacking strong sequence preference (31). Prior to sucrose-gradient centrifugation, lysates were treated with 1 U μL^-1^ of RNase I at different temperatures and reaction times (**Fig. 1F**). Polysome-collapse assays were then used to evaluate digestion efficiency based on conversion of polysomes into stable 80S monosomes (32). All digestion conditions reduced polysome abundance and increased the monosome (80S) peak in sucrose-gradient profiles, indicating efficient mRNA cleavage. These results further show that *Ectocarpus* monosome integrity is preserved following RNase I treatment.

Monosome fractions from each nuclease condition were collected and RFP libraries were prepared using a low-input protocol based on the usage of solid-phase reversible immobilization (SPRI) beads for purification of intermediates during library preparation (33). Ribo-seq libraries were sequenced to a depth of 29-47 M reads, and RNA libraries to 30**–**36 M reads. Data quality was assessed using riboWaltz and ORFik, which provide metrics for P-site offsets, reading-frame distribution, read length, and metagene footprint distribution across transcript features (34, 35).

Ribo-seq datasets generated using 1 U µL⁻¹ RNase I across all digestion temperatures and durations showed weak three-nucleotide periodicity, broad footprint length distributions, and reduced mRNA mapping relative to noncoding and intergenic regions (**Supplementary Fig. 1F–J**), consistent with incomplete nuclease digestion. The condition showing the strongest periodicity was RNase I digestion at 25 ⁰C for 60 min.

We therefore carried out RNase I digestion at 25 ⁰C for 60 min and increased RNase I concentration to 4.25 U µL⁻¹, then repeated library preparation and sequencing, using only male GA. Under these conditions, footprint lengths clustered at 28–31 nt, as expected for ribosome-protected fragments (**Fig. 1G**), and frame-0 P-site periodicity increased markedly (**Fig. 1H–J**). Metagene analysis showed strong enrichment of footprints within coding sequences compared to 5’ and 3’ untranslated regions (UTRs) (**Fig. 1K**). Footprint distribution indicated a ramp at the beginning of the ORFs and accumulation at stop codons, consistent with low elongation speed close to ORF start and ribosome pausing during translation termination, respectively (**Fig. 1K**). Elevated signal in 5′ UTRs, the lack of pronounced start codon peak in the metagene analysis (**Fig. 1K**), and mapping of reads to intergenic DNA (**Fig. 1L**) likely reflect annotation limitations in the current *Ectocarpus* genome assembly. Importantly, the majority of reads mapped to mRNA, a hallmark feature of translation (**Fig. 1L**).

Together, these results demonstrate that we successfully established a high-quality ribosome profiling protocol for brown algae. The strong frame bias, clear stop codon accumulation, and appropriate footprint size distribution indicate that our data are suitable for investigating translational regulation in *Ectocarpus* development.

#### Translational regulation distinguishes gene expression between sexes and contributes to developmental identity

To investigate the contribution of translational control to gene expression in *Ectocarpus* life cycle, we examined the relationship between mRNA abundance, ribosome occupancy and protein levels. For this purpose, we determined the proteome of the GA (male and female) and pSP (male) tissues by mass spectrometry LC/MS-MS (**Table S2**). We then estimated the correlation, using Spearman ρ, between mRNA abundance or ribosome occupancy to the abundance of proteins detected in each tissue (n=2400-2600 proteins) (**Supplementary Fig. 2A–C**). Independently of tissue type, protein abundance correlated better with ribosome occupancy than mRNA levels alone, as evidenced by higher Spearman correlation scores (Spearman ρ: Ribo-seq *vs* Protein=0.588±0.005, RNA-seq *vs* Protein=0.500±0.028; **Supplementary Fig. 2A–C**). These results highlight that protein levels are only partially determined by transcripts levels, with a significant proportion of regulation occurring at the post-transcriptional level. The strongest correlation was observed between mRNA levels and ribosome occupancy (ρ = 0.685 ± 0.036), consistent with both being derived from sequencing-based quantification, whereas protein abundance was measured by an independent mass spectrometry-based approach.

We therefore sought to identify the genes subject to translational control during *Ectocarpus* development. To this end, we employed deltaTE, which uses a DESeq2-based interaction model to test for changes in translation efficiency (TE) independent of variation in mRNA levels (36). Applying the deltaTE framework to the male versus female GA comparison, we categorized genes into regulatory classes based on their behavior across mRNA abundance and translation layers (**Fig. 2A, Table S3**). *Forwarded* genes (n = 3,235) are not subjected to translational control, displaying coordinated changes in both mRNA abundance and ribosome occupancy, i.e., translation is proportional to mRNA levels. Genes undergoing translational control spread among different categories according to the relationship with mRNA levels. *Buffered* genes (n = 569) exhibit compensatory translational regulation opposite to variation in mRNA abundance. *Exclusive* (n = 347) genes only show significant changes in ribosome occupancy (not mRNA levels) representing genes regulated exclusively by translation. *Intensified* (n = 87) genes positively or negatively amplify the signal of the mRNA levels by increasing or decreasing the number of ribosomes in the corresponding transcripts, respectively. Genes without differential expression were classified as *Unchanged* (n = 6565). All genes excluded during DESeq2 analysis due to lack of reliable Ribo-seq and RNA-seq read counts, or not meeting the combinatorial significance criteria required for deltaTE category assignment were assigned to the *Other* category (n = 7,468).

**Figure 2.**
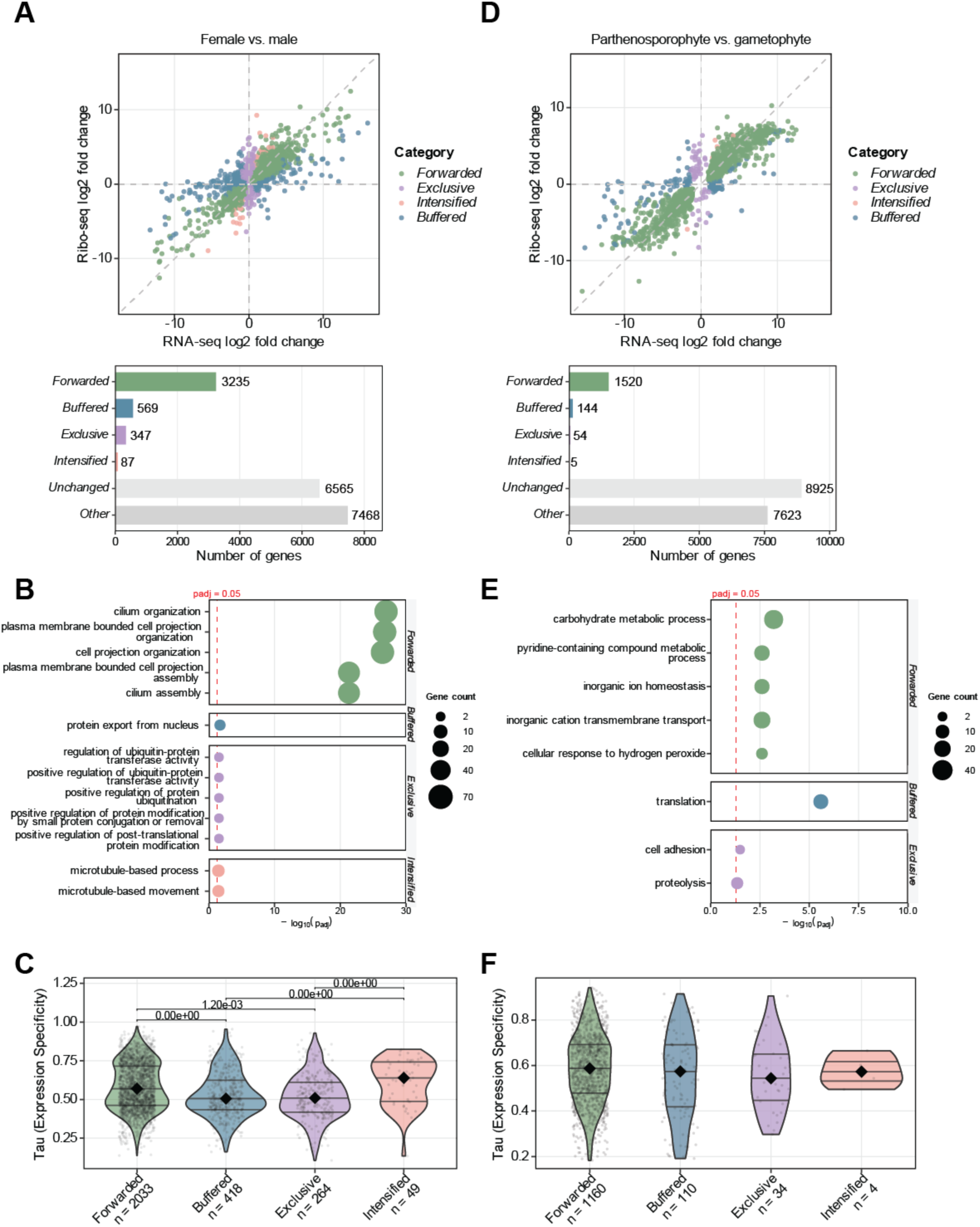
Translational regulation distinguishes gene expression between sexes and developmental stages. (A, **D)** DeltaTE analysis of female vs. male GA **(A)** or male GA vs pSP **(D)**. Upper graph: scatter plot of Ribo-seq vs RNA-seq log₂ fold changes, with genes colored by regulatory category (*Forwarded, Buffered, Exclusive, Intensified*). Dashed lines indicate the diagonal and quadrants. Lower graph: number of genes per category. **(B, E)** GO Biological Process enrichment of translationally regulated gene categories. Dot plot showing up to the top five significantly enriched GO terms (FDR < 0.05, Fisher’s exact test with Benjamini-Hochberg correction) per deltaTE category, in the female vs. male comparison **(B)** and in the GA vs pSP comparison **(E)**. Dot size indicates the number of significant genes annotated with each term. Only categories with at least one significant term are shown. **(C, F)** Tissue specificity (tau index) distributions of the genes with significant variation in expression in female vs male GA **(C)** or male GA vs pSP **(F)** samples. Genes are grouped by categories. Lower tau indicates broader expression. Sample sizes (n) are shown for each category. A permutation Kruskal-Wallis test (10,000 permutations) was used to assess overall group differences; significant comparisons were followed by pairwise permutation tests on median differences with Bonferroni correction.

Functional enrichment analysis (Gene Ontology, GO) revealed distinct roles for each regulatory category for specific biological processes (**Fig. 2B**, **Table S4–S5**). Comparing males and females, *Forwarded* genes were enriched for GO terms relating to cilium organization and assembly, and cell projection organization, which are both associated with flagellar function. These flagellar-related genes showed primarily female-biased gene expression. *Buffered* genes were associated with protein export from nucleus, whereas *Exclusive* genes were enriched for ubiquitin-related processes, however note there were only two genes under these categories. *Intensified* genes were similarly enriched in gamete or flagellar-related processes as *Forwarded* genes, with annotations related to dynein, axoneme, and kinesin; these also showed female-biased gene expression (**Table S4–S5**).

We also assessed the tissue expression range of the differentially expressed genes using tau (37, 38), a metric ranging from 0 (broadly expressed) to 1 (tissue-specific). Tau was previously determined for each *Ectocarpus* gene in (39) based on a comprehensive transcriptomic dataset spanning multiple tissue types and developmental stages. Genes subject to sex-specific translational regulation in the *Buffered* and *Exclusive* categories exhibited significantly lower Tau values than *Forwarded* genes (**Fig. 2C**), indicating that translational control preferentially targets genes with broad or multi-context expression rather than those with restricted, tissue-specific activity.

Comparison between the developmental stages GA and pSP using deltaTE analysis showed that *Forwarded* genes (n = 1,520) were the most prevalent category, followed by *Buffered* (n = 144), *Exclusive* (n = 54), and *Intensified* genes (n = 5) (**Fig. 2D, Table S6**). The markedly lower number of translationally regulated genes in the pSP versus mGA comparison, together with the higher proportion of *Unchanged* genes (8,925 vs. 6,565), indicates a greater overall similarity in gene expression at both the transcriptome and translatome levels between developmental stages than between sexes. The *Other* category comprised a comparable number of genes (n = 7,623).

GO analysis of biological processes showed that *Forwarded* genes were enriched in core metabolic and ion homeostasis functions (**Fig. 2E**, **Table S7–S8)** including both pSP- and GA-biased genes. *Buffered* genes were enriched for translation-related genes, notably 14 ribosomal proteins from both subunits, and displayed GA-biased expression at the Ribo-seq level. *Exclusive* genes were enriched for processes related to cell adhesion and proteolysis and showed predominantly pSP-biased translation. A CDC20 domain-containing protein (Ec-28_000800) within the proteolysis category overlapped with ubiquitin-associated *Exclusive* genes identified in the female versus male comparison. No significant enrichment was detected for *Intensified* genes.

Tau values did not differ significantly among gene categories in the GA versus pSP comparison (**Fig. 2F**), indicating similar tissue expression breadth between *Forwarded* genes and those subject to translational regulation (*Buffered, Exclusive, and Intensified*).

Together, these results demonstrate that translational regulation constitutes a critical layer in the control of gene expression in *Ectocarpus*, with distinct regulatory categories exhibiting specific functional specializations that ensure appropriate protein output across developmental stages and sexes.

#### Sequence features and selective translational control of developmental regulators in *Ectocarpus*

Having identified genes subject to translational control and their associated potential biological functions, we next examined sequence features that might underlie their regulation. Previous studies in diverse organisms have implicated codon usage, UTR length and UTR secondary structure as determinants of translational efficiency (12). We therefore systematically analyzed these features across regulatory categories.

We calculated the effective number of codons (ENC) for each gene as a measure of codon diversity, with lower values indicating stronger codon bias (40). Sex- and stage-related *Exclusive* and *Buffered* genes were the categories with significantly lower ENC values (**Fig. 3A-B**), indicating reduced codon diversity among translationally regulated transcripts. To assess whether this reduced diversity reflects codon optimization, we computed the Codon Adaptation Index (CAI) using ribosome occupancy-derived optimal codons as a reference (41), and GC content at synonymous third codon positions (GC3s) (40). As expected for a GC-rich genome in which all 18 optimal codons are GC-ending (**Supplementary Fig. 3A**), genes with low ENC values also exhibited high CAI and GC3s values, confirming that reduced codon diversity in *Ectocarpus* reflects preferential use of translationally optimal, GC-ending codons. Although CAI and GC3s differences across categories did not reach statistical significance in most comparisons (**Fig. C–F**), *Buffered* and *Exclusive* genes consistently showed higher CAI and GC3s values relative to *Forwarded* and *Intensified* genes. Together, these results indicate that translationally regulated genes in *Ectocarpus* are characterized by optimized codon usage, consistent with selection for translational efficiency and accuracy.

**Figure 3.**
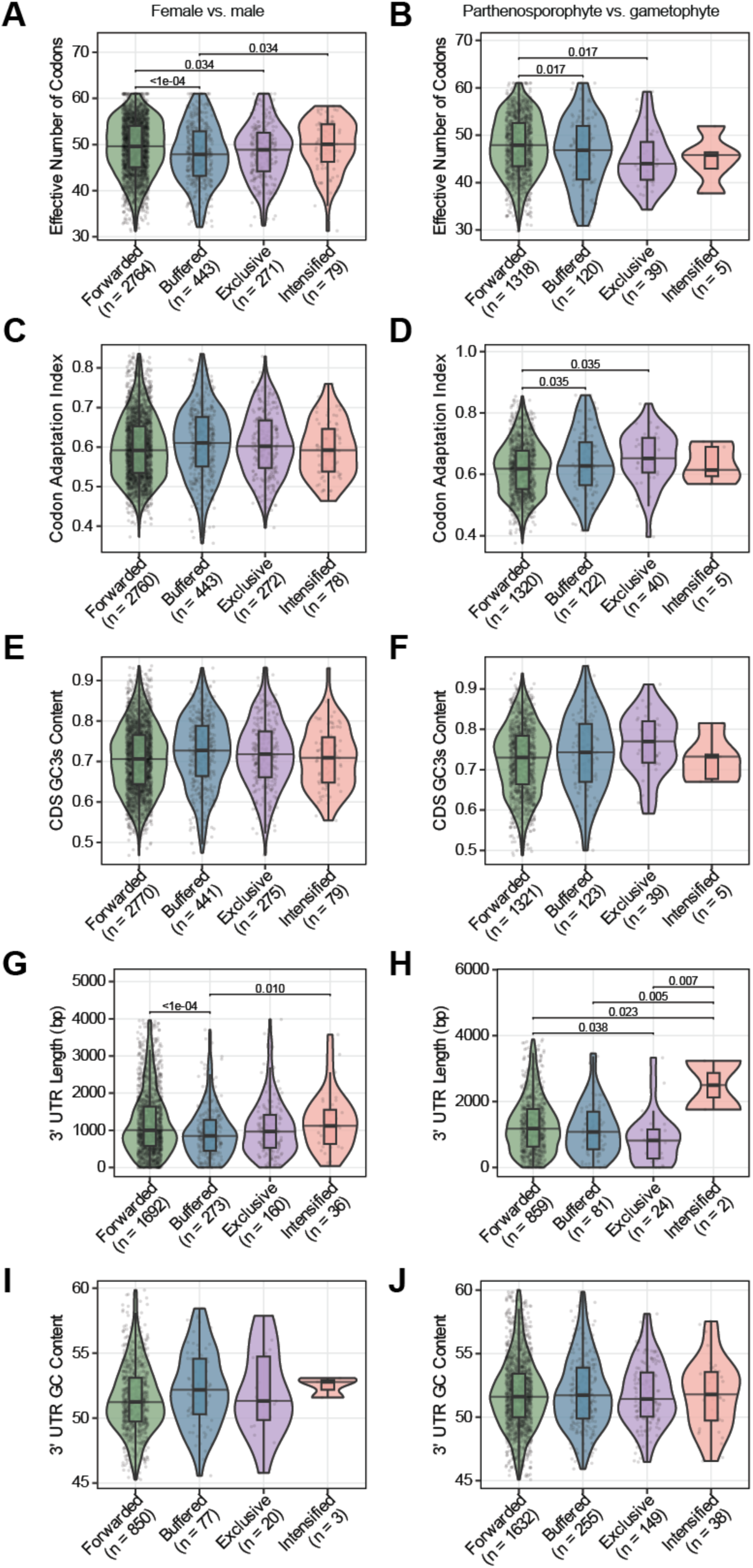
Sequence features of translationally regulated genes. Violin plots comparing sequence features across deltaTE regulatory categories for the female vs. male GA comparison (left column: A, C, E, G, I) and the pSP vs. male GA comparison (right column: B, D, F, H, J). Overlaid boxplots show median and interquartile range; outliers beyond 1.5× IQR were excluded for visualization. Sample sizes (n) are indicated for each category. A permutation F-test (10,000 permutations) was used to assess overall group differences. Significant features were followed by pairwise permutation tests on mean differences with FDR correction, and FDR-corrected p-values are shown above significant comparisons. Panels show **(A, B)** Effective Number of Codons (ENC), **(C, D)** Codon Adaptation Index (CAI), **(E, F)** GC content at synonymous third codon positions (GC3s), **(G, H)** 3′ UTR length, and **(I, J)** 3′ UTR GC content.

Among all categories, *Intensified* genes had longer 3′ UTRs, whereas sex-related *Buffered* and stage-related *Exclusive* genes possessed significantly shorter 3′ UTRs (**Fig. 3G-H**). 3′ UTR GC content did not differ significantly between regulatory categories in either comparison (**Fig. I–J**), suggesting that bulk nucleotide composition of the 3′ UTR does not distinguish translationally regulated genes in *Ectocarpus*.

Differences in 5′ UTR length or GC content were not examined due to the current limitations in 5′ UTR annotation for *Ectocarpus*.

We next took a closer look into genes with known developmental roles in *Ectocarpus* under translational control. OUROBOROS (*ORO*, Ec-14_005920), a TALE homeodomain transcription factor, functions as a master regulator of the gametophyte-to-sporophyte transition (42, 43). Loss-of-function mutations in ORO cause homeotic conversion of the sporophyte into a functional gametophyte, demonstrating its essential role in deploying the sporophyte developmental program (42, 43). *ORO* was classified as *Exclusive* in the deltaTE analysis of pSP vs. male GA, with significantly increased ribosome occupancy in pSP (log₂ fold-change TE = 2.17) but non-significant change in mRNA levels, although RNA-seq counts showed a trend toward higher expression in pSP with notable inter-replicate variability (**Table S6, Supplementary Fig. 3B**). The translational upregulation of *ORO* in pSP is consistent with its known role as a master regulator of the sporophyte developmental program (42, 43). Consistent with the properties associated with translational regulation, *ORO* also shows broad developmental expression (τ = 0.56), strong codon bias (ENC = 42.32), optimized codon usage (CAI = 0.70, GC3s = 0.80), high propensity for RNA secondary structure in the 3′ UTR (–ΔG = 12.1), and a 3′ UTR length (893 nt) comparable to the median for this gene category (**Fig. 3, Supplementary Fig. 3B**).

MALE INDUCER (*MIN*, Ec-13_001750) is a HMG-box transcription factor recently identified as the male sex-determining gene in *Ectocarpus* (44). This gene belongs to the *Forwarded* category in the deltaTE analysis between sexes. *MIN* expression was detected almost exclusively in male GA, with negligible RNA-seq and Ribo-seq signal in female GA (log₂ fold-change RNA = –11.75), consistent with its role as the male sex-determining gene (**Supplementary Fig. 3C**). In contrast to *ORO*, *MIN* exhibits properties characteristic of tissue-specific, transcriptionally regulated genes: high tissue specificity (τ = 0.77), weak codon bias (ENC = 53.97), less optimized codon usage (CAI = 0.44, GC3s = 0.53) and lower propensity for RNA secondary structure in the 3′ UTR (–ΔG = 10.7) (**Supplementary Fig. 3C**). However, its 3′ UTR (493 nt) is relatively short, falling at the lower end of the distribution observed for Forwarded genes (**Fig. 3, Supplementary Fig. 3C**).

The contrasting regulatory modes of *ORO* and *MIN* transcription factors imply that translational control is selectively deployed among developmental regulators. The broad expression but exclusive translational regulation of *ORO* is consistent with a model in which ORO protein levels may be rapidly modulated in response to life cycle cues, such as non-cell autonomous factors (42, 43), without requiring changes in mRNA abundance. The broad developmental expression and translational upregulation of *ORO* are consistent with a model in which translational control provides an additional layer of rapid, post-transcriptional tuning of ORO protein levels in response to life-cycle cues, complementing transcriptional regulation.

#### Genomic determinants of baseline translation efficiency in *Ectocarpus*

To investigate the genomic features that determine TE independently of sex or developmental stage, we pooled ribosome profiling and RNA-seq data from all three stages and compared genes with high versus low TE. Unlike the deltaTE approach, which identifies genes whose translation changes between conditions, this global TE analysis captures the baseline translational capacity encoded by gene sequences. For each gene, TE was calculated as the ratio of ribosome occupancy to mRNA abundance. Genes were then categorized into high-TE (upper 25th percentile) and low-TE (lower 25th percentile) groups based on their TE distribution (**Table S9**). High-TE genes are translated efficiently across all stages, whereas low-TE genes show poor translation. Importantly, these baseline TE categories are independent from the deltaTE regulatory classes (*Forwarded, Buffered, Exclusive, Intensified*), which reflect condition-specific changes in translation.

Codon usage analysis revealed marked differences between high- and low-TE genes. High-TE genes displayed significantly lower ENC values (**Fig. 4A**), indicative of more restricted codon usage. Consistent with codon optimization, these genes also exhibited higher CAI (**Fig. 4B**) and increased GC3s (**Fig. 4C**), demonstrating a preference for translationally optimal, GC-ending codons. The concordance of low ENC, high CAI, and elevated GC3s supports the conclusion that codon usage bias in *Ectocarpus* is shaped by selection for translational efficiency.

**Figure 4.**
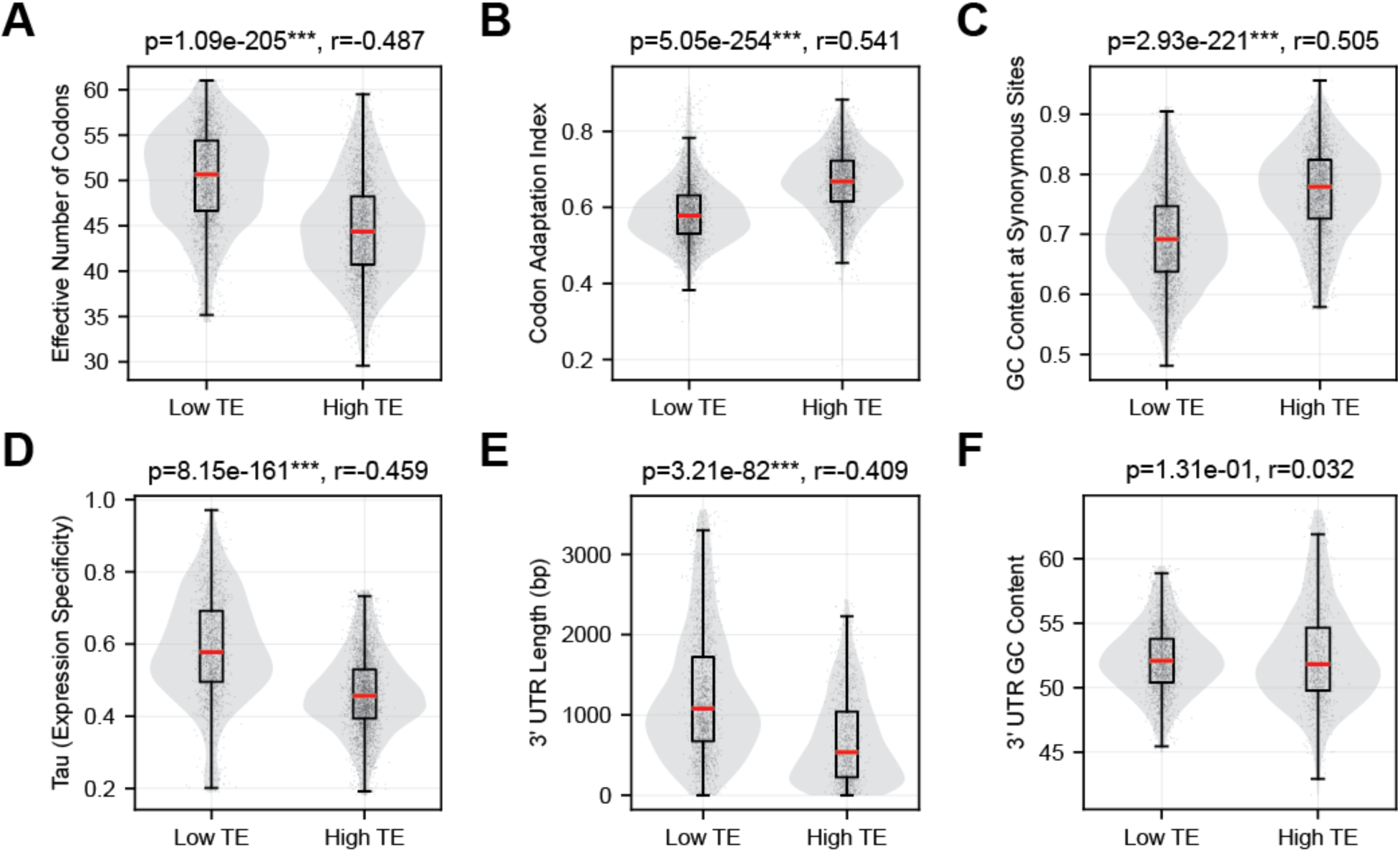
Sequence features of genes with high and low translation efficiency. For each gene, translation efficiency (TE) was calculated as the ratio of mean Ribo-seq to mean RNA-seq counts (DESeq2 normalized). Genes were classified as high TE (top 25th percentile) or low TE (bottom 25th percentile). Violin plots show distributions with boxplots indicating median and interquartile range. Statistical comparisons were performed using Wilcoxon rank-sum tests with Benjamini-Hochberg (FDR) correction: p-values and effect sizes (r) shown above each panel. Panels show **(A)** Effective Number of Codons (ENC), **(B)** Codon Adaptation Index (CAI, computed using Ribo-seq-derived optimal codons as reference), **(C)** GC content at synonymous third codon positions (GC3s), **(D)** tau expression specificity index, where values range from 0 (broadly expressed) to 1 (tissue-specific), **(E)** 3′ UTR length, and **(F)** 3′ UTR GC content.

In line with this, high-TE genes showed significantly lower tau values (**Fig. 4D**), indicating broader expression across developmental stages compared to low-TE genes. As broadly-expressed genes are required across multiple contexts, their sustained expression likely imposes stronger selective pressure for codon optimization to ensure efficient and accurate translation.

High-TE genes also possessed significantly shorter 3′ UTRs than low-TE genes (**Fig. 4E**), without a corresponding difference in GC content (**Fig. 4F**). Shorter 3′ UTRs may contribute to increased translation efficiency by limiting the presence of regulatory elements, such as miRNA target sites and RNA-binding protein motifs, that can modulate translation, mRNA localization and stability.

Together, these findings demonstrate that high translation efficiency in *Ectocarpus* is associated with specific genomic features: shorter UTRs, codon usage bias characterized by low ENC and high GC3s, and broader tissue transcription patterns.

## Discussion

### Establishment of high-quality ribosome profiling in brown algae

This study provides the first comprehensive ribosome-profiling analysis in brown algae and establishes *Ectocarpus* as a tractable system for investigating translational regulation in an understudied multicellular lineage. Our optimized protocol overcomes key technical challenges associated with brown algal physiology, including adaptation to marine osmotic conditions and high polysaccharide and polyphenol content, and yields data that meet or exceed quality standards reported in established model systems (13, 29, 33, 45, 46). The strong three-nucleotide periodicity, clear accumulation of reads at stop codons, and high phase scores demonstrate that ribosome profiling can be successfully applied to non-model marine organisms following appropriate optimization.

A key feature of the protocol was optimization of salt concentrations for ribosome isolation, with 200 mM NaCl required for GA, but 50 mM NaCl being sufficient for pSP. These stage-specific requirements highlight the importance of tailoring lysis conditions to cellular physiology across the life cycle and may inform adaptation of ribosome profiling to other marine algae or osmotically specialized organisms.

Efficient generation of periodic ribosome footprints required substantially higher RNase I activity than reported for yeast or mammalian systems. Optimal digestion in *Ectocarpus* (4.25 U μL⁻¹ lysate; ∼85 U μg⁻¹ RNA) corresponds to approximately 100–600-fold greater enzyme usage than in previously described protocols (33). This likely reflects increased resistance of endogenous RNA to nuclease digestion or inhibition of RNase I activity, and highlights the need for organism-specific optimization using polysome collapse assays when establishing ribosome profiling in new systems.

Data quality was further supported by the stronger correlation between ribosome occupancy and protein abundance than between mRNA and protein levels across all samples, consistent with observations in yeast, mammals, humans, and plants (47). The divergence between transcriptome-proteome and translatome-proteome correlations in *Ectocarpus* indicates that translational regulation substantially contributes to protein output in *Ectocarpus* and provides a quantitative basis for the regulatory phenomena described here.

We did not detect a pronounced start-codon ribosome accumulation peak, which in many systems reflects rate-limiting translation initiation. Because core translational machinery is highly conserved (48), this absence is unlikely to indicate fundamentally altered initiation dynamics. Instead, incomplete annotation of coding sequence start sites leading to misassignment of coding regions as 5′ UTR may obscure initiation signatures. In addition, brief delays between harvesting and flash-freezing could permit ribosome redistribution from initiation to elongation states, although prior studies more commonly report increased, rather than decreased, start-codon occupancy under such delays (49). Further refinement of genome annotation and rapid-harvest methodologies will therefore be important for resolving initiation dynamics in brown algae.

Overall, the ribosome-profiling framework established here should be broadly transferable to other brown algae and potentially to diverse marine organisms with similar biochemical and osmotic constraints, provided that lysis ionic conditions and nuclease digestion parameters are empirically optimized.

#### Translational control fine-tunes developmental gene expression in *Ectocarpus*

Our differential expression analyses across sexes and developmental stages indicate that translational regulation contributes to the definition of different developmental programs associated with *Ectocarpus* life cycle. The importance of translational regulation to the establishment of the different proteomes is further evidenced by the higher correlation between translatomes and proteomes than between transcriptomes and proteomes of the different tissues here studied. Interestingly, translational regulation had a stronger contribution to sex-specific tissue transitions than to stage differentiation (GA vs pSP). The broadly similar gene expression programs of GA and pSP are consistent with previous transcriptomic and epigenetic studies (26, 50), and supported by *Ectocarpus* having a weak dimorphic life cycle (51), meaning the GA and (p)SP stages are similar in morphology. There are other brown algae with stronger morphological differences between the GA and (p)SP stages such as *Saccorhiza polyschides*, *Macrocystis pyrifera*, or *Saccharina latissima* (51). In these species translational control may have a higher contribution to the stage differentiation programs.

Interestingly, we found that *Forwarded* and *Intensified* genes in the sex comparison were enriched for gamete-related functions. Previous studies have reported a higher density of gametangia in male *Ectocarpus* GA compared to females (27), which would suggest that these enriched gamete-related genes are likely to be male-biased in their expression at both the transcriptional and translational levels. Surprisingly, however, these genes instead exhibited a clear female-biased expression pattern. This unexpected result suggests that factors beyond visible morphological differences may be at play. We propose that underlying metabolic or physiological differences, not readily apparent from morphology alone, may contribute to the observed female bias in gamete-related gene expression (26, 44, 52). By contrast, the limited transcriptomic and epigenetic differences previously reported between gametophyte and partheno-sporophyte generations align with the reduced translational divergence detected here (26, 50).

Genes under translational selection were classified into *Buffered*, *Exclusive*, and *Intensified* categories, revealing a clear predominance of translational buffering. In this mode, changes in ribosome occupancy counteract variation in transcript levels, thereby stabilizing protein output. Although only recently recognized, translational buffering is increasingly understood as a fundamental layer of gene regulation across different biological systems (53). It has been shown to preserve protein levels across divergent genetic backgrounds and stress conditions in yeast (13, 47, 54) to constrain protein variability across tissues in mammals (55).

In our pSP vs GA comparison, *Buffered* genes were significantly enriched for the GO term “translation,” including many components of the ribosomal machinery. This pattern aligns with previous observations that genes encoding subunits of macromolecular complexes, especially ribosomal proteins, are preferentially subject to buffering, likely reflecting the need to maintain precise stoichiometric balance (55). Taken together, our results extend this conserved regulatory principle to brown algae, highlighting translational buffering as a central and evolutionarily conserved mechanism for maintaining protein homeostasis, particularly for core cellular machinery where dosage imbalance would be most detrimental.

The molecular mechanisms underlying translational buffering in *Ectocarpus* remain to be elucidated. In other systems, buffering has been linked to specific 5′ UTR features, such as upstream open reading frames and RNA secondary structures, as well as to the action of RNA-binding proteins and RNA modifications (12). Whether similar mechanisms operate here is an open question. Identifying the relevant cis-elements and trans-acting factors will therefore be an important goal for future work.

From a broader perspective, our results complement existing knowledge of transcriptional regulation by showing that gene expression differences between sexes and life stages are not solely determined at the mRNA level. While transcriptional programs likely establish the primary framework of sex- and stage-specific expression, translational regulation adds an additional layer of control that can buffer, reinforce, or uncouple these differences. In particular, the observed enrichment of metabolic and physiological functions among differentially regulated genes may reflect underlying functional divergence between tissues that is not fully captured by morphology alone.

Beyond condition-specific translational regulation, our analysis reveals that intrinsic sequence features are strong determinants of baseline translational output. Genes with constitutively high translation efficiency are characterized by optimized codon usage - low ENC, high CAI, and high GC3s - shorter 3′ UTRs, and broader expression across developmental stages. In *Ectocarpus*, all 18 optimal codons identified from the ribosome occupancy-derived reference set are GC-ending, consistent with the GC-rich genome driving mutational bias toward GC-ending codons. The convergence of codon optimization, compact 3′ UTRs, and broad expression among high-TE genes suggests that intrinsic sequence properties predispose certain transcripts to efficient translation. Conversely, low-TE genes exhibit weaker codon bias, longer 3′ UTRs, and more tissue-restricted expression. This distinction implies that the genomic architecture of a transcript is a major determinant of its baseline translational output, with codon optimization and UTR length acting as complementary features that together shape translational capacity independently of condition-specific regulatory mechanisms.

#### Homeostatic and developmental roles of translational control in *Ectocarpus*

An important consideration in interpreting our results is that the experimental design captures steady-state adult tissues from fully differentiated gametophytes and partheno-sporophytes rather than cells undergoing active developmental transitions. The translatome described here therefore reflects maintenance of established tissue identity rather than the dynamic processes that generate it. In systems profiled during developmental transitions, such as yeast meiosis (45), *Drosophila* oocyte-to-embryo transition (18), or *Arabidopsis* early seed maturation (19), substantially more extensive translational reprogramming is typically observed. Our analysis thus likely underestimates the full extent of translational regulation during *Ectocarpus* development.

Nevertheless, the detection of substantial translational regulation in steady-state tissues underscores its importance as a homeostatic mechanism that maintains appropriate protein levels once developmental programs are established. The predominance of *Buffered* genes is consistent with this interpretation: in differentiated tissues, translational buffering may dampen transcriptional variability and preserve the precise protein stoichiometries required for stable cellular function rather than driving large-scale proteome remodeling.

In summary, this study establishes ribosome profiling in a brown alga and provides the first comprehensive characterization of translational regulation in an independently evolved multicellular eukaryotic lineage. Translational buffering emerges as the predominant post-transcriptional regulatory mechanism in *Ectocarpus*, consistent with its conservation across fungi and animals. Our study positions brown algae as a key evolutionary system for uncovering fundamental principles of post-transcriptional gene regulation in multicellular eukaryotes.

## Material and Methods

### Algal culture and sample preparation

*Ectocarpus* strains were maintained and cultivated in half-strength Provasoli-enriched autoclaved seawater at 15°C. Gametophyte clones were initially grown on glass coverslips in 7-8 mL polystyrene Petri dishes under low-light conditions (20 μmol photons m⁻² s⁻¹, 12 h:12 h light:dark cycle) for 3 weeks. Subsequently, algae were transferred to 100 mL polystyrene Petri dishes and grown under normal-light conditions for 3-4 weeks, with approximately 10 individuals per plate. Male partheno-sporophytes were propagated asexually by cutting basal tissue of adult pSPs into small fragments with a sterile scalpel and allowing regeneration for 4 weeks under normal-light conditions, with 15-20 pSPs per 100 mL dish.

Adult algae were harvested by filtering through a metal sieve with 71 μm pore size, briefly dried in tissue paper, immediately flash-frozen in liquid nitrogen, and stored at −80°C until used. Three biological replicates were collected for each developmental stage (female GA, male GA and male pSP), where each replicate consisted of tissue pooled from 10–15 culture plates. The same biological replicates were used for Ribo-seq and RNA-seq. A separate tissue harvest with the same growth conditions was performed for proteomics.

#### Tissue Lysis

Male gametophyte and sporophytes and female and male gametophyte were harvested 3-4 weeks following culture in 10-15 100 ml plates. The collected algal tissues were ground in liquid nitrogen using a mortar and pestle until a fine powder was obtained. 100 mg of frozen powder was transferred to 2.0 mL Eppendorf tubes without thawing. Lysis buffer (250 μL) containing 100 mM Tris-HCl pH 7.4, 1% sodium deoxycholate, 2% polyoxyethylene-10-tridecylether, 2% Plant RNA Isolation Aid (Invitrogen), 1 mM DTT, and 100 μg/mL cycloheximide was prepared with three different salt concentrations: low salt (50 mM NaCl, 5 mM MgCl₂), medium salt (200 mM NaCl, 20 mM MgCl₂), or high salt (700 mM NaCl, 70 mM MgCl₂). Lysates were vortexed thoroughly and mixed on a rotator for 10 min, then centrifuged at 4,000 × g for 5 min. To obtain cleared lysates, the supernatants were transferred to new tubes and centrifuged at maximum speed (>20,000 × g) for 10 min.

#### Polysome profiling (density gradient ultracentrifugation)

Sucrose density gradients (10-50% w/v) were prepared in ultracentrifuge tubes (Seton, S7022) using a Gradient Master (Biocomp). Gradient solutions contained 100 mM Tris-HCl pH 7.4, 50 μg/mL cycloheximide, and the low, medium or high salt concentrations described in the Tissue lysis section. 250 μL of cleared lysate was loaded onto each (low, medium and high salt) gradient and centrifuged using a SW55Ti rotor in a Beckmann Optima XPN-90 ultracentrifuge at 45,000 rpm (245,418.9 × g) for 1 h 15 min at 4°C. Absorbance at 260 nm was measured throughout the gradient using a piston gradient fractionator (Biocomp) and 12x 400 μl fractions were collected for further analysis.

RNA was isolated from the fractions by acid phenol-chloroform extraction. Samples were diluted with 0.5 v/v nuclease-free water and mixed with equal volume of Acid-Phenol:Chloroform, pH 4.5 (with IAA, 125:24:1) (Thermo Fisher). RNA was precipitated using 1/10 volume of 3 M sodium acetate pH 5.5 (Thermo Fisher), 2 uL linear acrylamide, and 0.8 volumes isopropanol. Washing was done with cold 80% ethanol, and the air-dried pellet was resuspended in 15 uL of nuclease-free water.

Purified RNA was mixed with 1x Green GoTaq buffer (Promega) as loading dye, separated on a 1.2 % agarose gel stained with Midori Xtra (Nippon Genetics Europe) according to manufacturer instructions, and analyzed for 18S and 28S rRNA bands. Experiments were carried out three times with similar results.

Polysome collapse assay was done by carrying out polysome profiling assays after treatment with different RNase I (Ambion) concentrations (1 or 4.25 U/uL RNase), or digestion times (30 min, 45 min, 60 min, 18 h), or temperature (4 or 25 ⁰C).

#### Ribosome profiling library preparation

Ribosome profiling was performed as described in Meindl *et al.* (33) with slight modifications. Cleared lysates obtained following tissue lysis were treated with 1 U/µL RNase I (Ambion) at 25 ⁰C for 30 min, 45 min, and 1 h, or at 4 ⁰C for 18 h, or with 4.25 U/uL RNase I (Ambion) at 25 ⁰C for 1 h. Following nuclease digestion, monosomes were purified using sucrose density gradient centrifugation as described above (polysome profiling section) by collecting the fraction of the sucrose density gradient corresponding to the 80S monosomes. RNA was purified by acid phenol-chloroform extraction. Samples were diluted with 0.5 v/v nuclease-free water and mixed with equal volume of Acid-Phenol:Chloroform, pH 4.5 (with IAA, 125:24:1) (Thermo Fisher). RNA was precipitated using 1/10 volume of 3 M sodium acetate pH 5.5 (Thermo Fisher), 2 uL linear acrylamide, and 0.8 volumes isopropanol. Washing was done with cold 80% ethanol, and the air-dried pellet was resuspended in 20 uL of nuclease-free water. Purified RNA was mixed with 1x formamide loading buffer (50 mM EDTA-NaOH pH=8.0, 0.05% (w/v) bromophenol blue in formamide), and separated on 17% TBE-urea gels and visualized using SYBR Gold (ThermoFisher) staining. RNA oligonucleotides Marker-27nt and Marker-30nt (**Table S10**) were used as size markers to guide the excision of the 26-32 nt ribosomal footprint band. RNA was extracted from the gel piece by incubating with 400 µL extraction buffer (300 mM sodium acetate pH 5.5, 1 mM EDTA, 0.25% SDS, SUPERase In (Invitrogen) overnight, rotating at 4 ⁰C. RNA was precipitated using 2 volumes of ethanol and 2.5 µL of linear acrylamide for 3 h at −80 ⁰C, pelleted for 30 min at 4 ⁰C using max speed in a bench-top centrifuge, then resuspended in 17 µL of nuclease-free water. Terminal phosphates from RNase I digestion were removed from the isolated ribosome footprints using T4 PNK (New England Biolabs) according Meindl et al. (33). 3′ adapter ligation was performed using T4 RNA ligase II truncated KQ (New England Biolabs) with pre-adenylated rApp-L7 adapters (constructed by using Mth RNA ligase/5’ DNA Adenylation Kit (NEB) on on 5’ phosphorylated rApp-L7 according to manufacturer’s instructions. Ribosomal RNA depletion was carried out using subtractive hybridization with custom made siPOOL oligonucleotides (siTOOLs Biotech) complementary to *Ectocarpus* rRNA sequences following the manufacturer’s instructions. The ligated ribosome footprint RNA was isolated using Dynabeads MyOne Silane beads (ThermoFisher). and reverse transcribed using Superscript III (Thermo Fisher) and 20 pmol of P7 RT oligonucleotide according to the manufacturer’s protocol. Hydrolysis of RNA was done as in Meindl et al. (33) – by incubating with 1 M NaOH at 98 ⁰C for 20 min, then neutralized with 1M HEPES-NaOH pH 7.3. cDNA was purified using Dynabeads MyONE Silane beads (Thermo Fisher) as described above. cDNA was circularized using CircLigase (Lucigen) and purified using MyONE Silane beads (Thermo Fisher).

Initial library amplification was performed using 2× KAPA HiFi HotStart ReadyMix (Roche) for 6 cycles with primers P5Solexa_s and P7Solexa_s (**Table S10**). ProNex Size-selective Purification beads (Promega) were used at 1:2.95 v/v ratio to purify the PCR product. Scouting PCRs with 6-11 cycles were performed and visualized on agarose gels to determine optimal cycle number for library amplification. Final library amplification PCR included multiplexing using NEBNext Index Primers for Illumina (New England Biolabs, **Table S10**). PCR products were purified on 3% agarose cassettes selecting fragments of 130-208 bp length using the “range” setting on a Pippin Prep system (Sage Science). Final quality analysis was performed on a Bioanalyzer 2100 Expert High Sensitivity DNA Assay. Libraries were sequenced at a depth of 29 - 47 million reads single-end 50 bp reads on a NovaSeq X Plus platform (Novogene).

#### RNA-seq library preparation

Total RNA was extracted from all biological samples using TRIzol LS reagent (Invitrogen) according to the manufacturer’s protocol. RNA quality was assessed using a Bioanalyzer 2100. RNA-seq libraries were prepared using the NEBNext® Ultra II Directional RNA Library Prep Kit for Illumina® (Illumina) and NEBNext Poly(A) mRNA Magnetic Isolation Module according to the manufacturer’s instructions. Libraries were sequenced on a NextSeq2000 platform (Genome Center, MPI Biology Tübingen) to generate approximately 30 million 150 bp paired-end reads per sample.

#### Data processing and quality control

Ribo-seq and RNA-seq reads were processed using nf-core/riboseq (56) version 1.2.0 and nf-core/rnaseq (57) version 3.19.0 pipelines, respectively. In both pipelines, adapters were trimmed and reads were quality-filtered using Trim Galore! (58). In nf-core/riboseq, UMI extraction and deduplication was done using UMI-tools (59), and reads mapping to ribosomal RNA were removed using SortMeRNA (60). For both pipelines, high-quality reads were aligned to the *Ectocarpus* genome Ec32_v5 (61). STAR aligner (62) was used with the parameters differing from the nf-core defaults: *--alignIntronMax 50000 --outFilterMismatchNmax 1 --alignEndsType Local* for Ribo-seq data, and *--alignIntronMax 50000* for RNA-seq data. Read counts were quantified at the gene level using Salmon (63).

To perform comprehensive quality control analyses, ribosome profiling reads were analysed with ORFik (35) and riboWaltz (34). P-site offset determination was performed using riboWaltz’s psite function with automatic extremity detection and a flanking region of 6 nucleotides. riboWaltz was additionally used to assess trinucleotide periodicity by computing the percentage of P-sites mapping to each reading frame within the CDS. To assess data quality at the transcript level, proportion of P-site reads in the correct reading frame (frame 0) per transcript were calculated using riboWaltz’s frame_psite_length output. Metagene profiles were generated by extracting ribosome footprint coverage across scaled 5′ UTR, CDS, and 3′ UTR regions for each transcript using ORFik. To enable averaging across transcripts with different expression levels, coverage values were Z-score normalized within each transcript and condition by subtracting the mean and dividing by the standard deviation of coverage across all positions. Normalized profiles were then averaged across all transcripts per condition. ORFik was also used for quantifying read mapping across genomic features. P-site-shifted reads were used for all downstream analyses.

#### Sample preparation for mass spectrometry

Proteome extraction was performed using a modified protocol adapted from Ritter et al. (64). Briefly, 100 mg of frozen algal tissue was ground in liquid nitrogen for 20 min in triplicates. Algal powder was resuspended in 2.5 mL of extraction buffer [(1.5 % PVP40, 0.7 M sucrose, 100 mM KCl, 100 mM Tris-HCl pH 7.5, 10 mM EDTA, 0.5 % CHAPS, 1x cOmplete Protease Inhibitor Cocktail (Roche)] and gently rotated at 4°C for 10 min. An equal volume (2.5 mL) of Tris-HCl pH 7.5 saturated phenol was added, and the mixture was homogenized for 10 min at 4°C on a shaker. The algal lysate was centrifuged at 10,000 × g for 20 min to achieve clear phase separation. The upper phenol phase was removed and extracted with 2.5 mL of extraction buffer, rotated for 3 min, vortexed, and centrifuged at 10,000 × g for 20 min. The upper phenol phase was retrieved and split into two tubes. Proteins in the phenol phase were precipitated with five volumes of 0.1 M ammonium acetate dissolved in methanol and incubated at −20°C for 3 h or overnight. The extract was centrifuged at 10,000 × g for 20 min, and the supernatant discarded. The protein pellet was rinsed in ammonium acetate (0.1 M in methanol) for 20 min at −20°C, and then washed four times in four volumes of 80% ice-cold acetone by inverting the tube to dislodge the pellet and centrifuging at 16,000 × g for 5 min. Each protein pellet was dissolved by pipetting up and down in 50 μL 1x SDS-PAGE loading buffer (100 μL per sample total), and heated to 80°C with shaking at 1000 rpm for 15 min.

#### In-gel digestion of proteins and mass spectrometry

In-gel digestion and mass spectrometry were done at the Proteome Center Tübingen. Proteins were loaded and run on a NuPAGE 12% Bis-Tris SDS-PAGE Gel (Thermo Fisher Scientific) and stained with colloidal Coomassie using the ReadyBlue Protein Gel Stain (Merck Millipore). Gel sections containing proteins were excised, and cut into smaller parts. For in-gel digestion of proteins, gel pieces were destained by washing three times with 5 mM ammonium bicarbonate (ABC) in acetonitrile (ACN) (1:1, v/v) for 20min. After a dehydration step with 100% ACN for 10min, disulfide bonds were reduced with 10 mM dithiothreitol (DTT) in 20 mM ABC for 45 min at 56°C, and thiol groups of cysteine residues were prevented from reoxidation by carbamidomethylation with 55 mM iodoacetamide (IAA) in 20mM ABC for 45 min in the dark. Gel pieces were then washed two times with 5 mM ABC in ACN (1:1, v/v) for 20min and dehydrated with 100% ACN for 15min. After evaporation of the liquid in a vacuum centrifuge for 10 min, gel pieces were soaked in a solution of 12.5 ng/µl sequencing grade trypsin (Promega) in 20 mM ABC, pH 8.0 for 10 min at room temperature (RT), and then covered with 20 mM ABC. After in-gel digestion of proteins at 37°C overnight, peptides were extracted in three consecutive steps with different extraction buffers for 30 min: first 3% (v/v) trifluoroacetic acid (TFA) in 30% (v/v) ACN was added, followed by 0.5% (v/v) formic acid (FA) in 80% (v/v) ACN, and finally by 100% ACN. ACN was evaporated from pooled supernatants by vacuum centrifugation. In the course of the digestion protocol all incubation steps were carried out under shaking.

Desalted peptides were analyzed on a Vanquish Neo nano-UHPLC coupled to an Orbitrap Exploris 480 mass spectrometer through a nano-electrospray ion source (all Thermo Scientific). Estimated amount of 0.25 µg peptides were loaded onto a 20 cm HPLC column with 75 μm inner diameter (CoAnn Technologies, ICT36007508F-50) in-house packed with 1.9 μm ReproSil-Pur C18-AQ silica beads (Dr. Maisch HPLC GmbH, r119.aq.) under pressure control (1000 bar). Elution was performed with a 45 min segmented gradient of 5-55% of HPLC solvent B (80% acetonitrile in 0.1% formic acid) at a flow rate of 300 nl/min and a constant temperature of 40°C.

#### Mass spectrometry data processing

MS data were searched against a target-decoy database of *Ectocarpus* (61) and commonly observed contaminants using the Andromeda search engine integrated into the MaxQuant software (v2.2.0.0) (65). Search parameters were kept to default. In addition, the iBAQ (Intensity Based Absolute Quantification) and LFQ (Label-Free Quantification) algorithms were enabled, as was the “match between runs” option (66).

#### Multi-omics correlation analysis

To assess the relationship between mRNA, ribosome occupancy, and protein levels, Spearman correlations were calculated between RNA-seq (TPM), Ribo-seq (TPM), and proteomics (iBAQ) data for genes detected across all three platforms within each developmental stage. Hexbin density plots were used to visualize pairwise correlations. For multi-omics correlations, TPM was used for RNA-seq and Ribo-seq to enable direct comparison with iBAQ-normalized protein abundances, as both TPM and iBAQ provide length-normalized estimates of relative molecular abundance within a sample. DESeq2-normalized counts, which do not correct for gene length, were used for all differential expression and translational efficiency analyses, where within-gene comparisons across conditions make length normalization unnecessary.

#### Differential expression and translational efficiency analysis

Differential translational efficiency analysis was performed using the deltaTE approach (36), which is based on DESeq2. Transcript-level counts were aggregated to gene level (by summing counts across transcripts of the same gene) prior to DESeq2 modeling. Count matrices for ribosome profiling (Ribo-seq) and RNA-seq were combined, and the following interaction model was fitted:

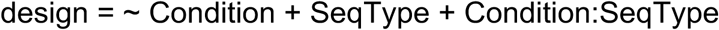

where Condition represents the biological comparison (e.g., female vs. male GA), SeqType distinguishes Ribo-seq from RNA-seq, and the interaction term Condition:SeqType captures differential translation efficiency (ΔTE). Genes with a significant interaction term (adjusted p-value < 0.05) were considered to have differential translational efficiency. Following the deltaTE methodology, differentially expressed genes were categorized into six groups based on the significance of changes in RNA-seq (padj_RNA), Ribo-seq (padj_Ribo), and TE (padj_TE):

*Forwarded*: genes showing significant changes in both RNA and Ribo-seq in the same direction, but no significant change in TE (padj_TE > 0.05, padj_Ribo < 0.05, padj_RNA < 0.05)

*Exclusive*: genes showing significant change in Ribo-seq but not RNA-seq, with significant ΔTE (padj_TE < 0.05, padj_Ribo < 0.05, padj_RNA > 0.05)

*Intensified*: genes showing significant changes in RNA, Ribo-seq, and TE in the same direction (padj_TE < 0.05, padj_Ribo < 0.05, padj_RNA < 0.05, ΔTE and ΔRNA show the same directional change)

*Buffered*: genes showing significant changes in RNA and TE in opposite directions, resulting in no change or reduced change in Ribo-seq (padj_TE < 0.05, with ΔTE and ΔRNA having opposite signs; or padj_TE < 0.05, padj_Ribo > 0.05, padj_RNA < 0.05)

*Unchanged/Housekeeping*: genes showing no significant changes in RNA, Ribo-seq, or TE (padj_TE > 0.05, padj_Ribo > 0.05, padj_RNA > 0.05)

*Other*: all remaining genes, excluded by DESeq2’s independent filtering of lowly expressed genes or those having outlier counts, or those not meeting combinatorial significance criteria required for deltaTE category assignment.

#### Functional enrichment analysis

Gene Ontology (GO) enrichment analysis was performed using topGO (67) with the “classic” algorithm and Fisher’s exact test. GO annotations for *Ectocarpus* genes were obtained from InterProScan (68) results mapped to GO terms, which was previously done in (39)Background gene sets included all genes in the master deltaTE table. Enriched terms with padj < 0.05 were considered significant.

#### Tissue specificity analysis

Tau values were obtained from (39) calculated across developmental stages and tissues. Where multiple transcripts mapped to the same gene, the maximum tau value was used. Differences in tau across regulatory categories were assessed using a permutation Kruskal-Wallis test (10,000 permutations), where the observed H-statistic was compared against a null distribution generated by randomly permuting category labels. When the global permutation test was significant (p < 0.05), pairwise comparisons were performed by permuting the difference in median tau between each pair of categories (10,000 permutations), with Bonferroni correction applied to the resulting p-values.

#### Codon Adaptation Index (CAI) calculation

The Codon Adaptation Index (CAI) was computed following the Sharp and Li (41) method using ribosome occupancy as a measure of gene expression. A reference set of optimal codons was determined from the top 10% of genes ranked by mean Ribo-seq normalized counts across all samples. For each amino acid with synonymous codons, the relative synonymous codon usage (RSCU) was calculated from this reference set, and the relative adaptiveness (w) of each codon was defined as the ratio of its RSCU to the maximum RSCU within the synonymous family. Gene-level CAI was computed as the geometric mean of the relative adaptiveness values across all codons in the coding sequence, excluding methionine, tryptophan, and stop codons.

#### Codon usage, UTR length and GC content analysis

Gene-specific codon usage metrics were calculated using CodonW (http://codonw.sourceforge.net/). For each protein-coding gene, the effective number of codons (ENC) and GC content at synonymous third codon positions (GC3s) were computed from the longest coding sequence. 3′ UTR sequences were extracted from the genome annotation, and UTR lengths and GC content were calculated for each principal transcript. Statistical comparisons of sequence features across deltaTE regulatory categories were performed using permutation F-tests (10,000 permutations). When the overall test was significant (p < 0.05), pairwise comparisons were performed using permutation tests on mean differences with FDR (Benjamini-Hochberg) correction. For the global TE analysis (high vs low TE groups), Wilcoxon rank-sum tests with Bonferroni correction were used, and effect sizes were reported as rank-biserial correlation (r).

#### RNA secondary structure prediction

5′ and 3′ UTR sequences of *ORO* (Ec-14_005920) and *MIN* (Ec-13_001750) were extracted from the *Ectocarpus* genome annotation. The annotation of *ORO* was updated using information from the RNA- and Ribo-seq data generated in this study (**Table S11**).

#### Global translation efficiency analysis

For each gene, TE was calculated as the ratio of mean ribosome occupancy to mean mRNA abundance across all samples, using DESeq2-normalized counts. For each TE group, we extracted the following genomic features: 3′ UTR lengths and GC content, effective number of codons (ENC), and GC content at third codon positions (GC3s). Statistical comparisons between high and low TE groups were performed using Wilcoxon rank-sum tests with FDR correction for multiple testing.

#### Gene-level coverage visualization

For individual gene coverage figures, P-site-shifted Ribo-seq read counts and RNA-seq read counts were extracted per nucleotide position across the transcript using pysam. RNA secondary structure propensity was predicted using a sliding window approach with RNAfold from the ViennaRNA package (69), computing the negative minimum free energy (–ΔG) in 40-nt windows with a 20-nt step across the transcript sequence. Coverage tracks for each condition were plotted alongside the gene model and RNA structure profile.

#### Statistical analysis and visualization

All statistical analyses were performed in R (version 4.3.0). For differential expression, multiple testing correction was applied using the Benjamini-Hochberg method. Pairwise comparisons of genomic features across regulatory categories were analyzed with Bonferroni correction. Visualizations were generated using ggplot2 (70). Volcano plots, heatmaps, and metaplots were generated using custom R scripts. All p-values reported are adjusted p-values unless otherwise stated.

## Code and data availability

All analysis scripts are available on Github at jiaxuanleong/translation_ectocarpus. Raw sequencing data have been deposited in NCBI BioProject with the accession number PRJNA1430186. Mass spectrometry data have been deposited in the PRIDE database with the project accession PXD075612. Processed count matrices and analysis results are available as **Supplementary Tables S1-11**.

## Author contributions

SMC and CI conceived and designed the experiments and supervised JXL; SMC acquired funding and supervised the project. JXL performed experiments with help of CI. JJBR and FH curated the data. JXL performed data analysis and visualization with support from FH, JBR and CI. JXL wrote the original draft with support from CI and SMC. SMC and CI reviewed and edited the manuscript.

## Acknowledgments

We thank the Genome Center MPI Biology Tübingen for sequencing support, the Proteome Center University of Tübingen for mass spectrometry support, and members of the Department of Algal Development and Evolution especially Jaruwatana Sodai Lotharukpong, Rita A. Batista, and Pélagie Ratchinski for helpful discussions. JXL thanks IMPRS ‘From Molecules to Organisms’.

## Supplementary Data

**Table S1.** *Ectocarpus* strains and culture conditions used in this study.

**Table S2.** Mass spectrometry-based protein quantification (iBAQ) across developmental stages.

**Table S3.** DeltaTE analysis results for the female vs. male gametophyte comparison.

**Table S4.** GO Biological Process enrichment analysis for deltaTE categories in the female vs. male gametophyte comparison.

**Table S5.** Genes associated with enriched GO terms in the female vs. male gametophyte comparison.

**Table S6.** DeltaTE analysis results for the parthenosporophyte vs. male gametophyte comparison.

**Table S7.** GO Biological Process enrichment analysis for deltaTE categories in the parthenosporophyte vs. male gametophyte comparison.

**Table S8.** Genes associated with enriched GO terms in the parthenosporophyte vs. male gametophyte comparison.

**Table S9.** Global translation efficiency values and expression data for all genes across developmental stages.

**Table S10.** Oligonucleotide sequences used for ribosome profiling library preparation.

**Table S11.** Updated 5′ and 3′ UTR sequences for *OUROBOROS* (Ec-14_005920).

**Supplementary Figure 1.**
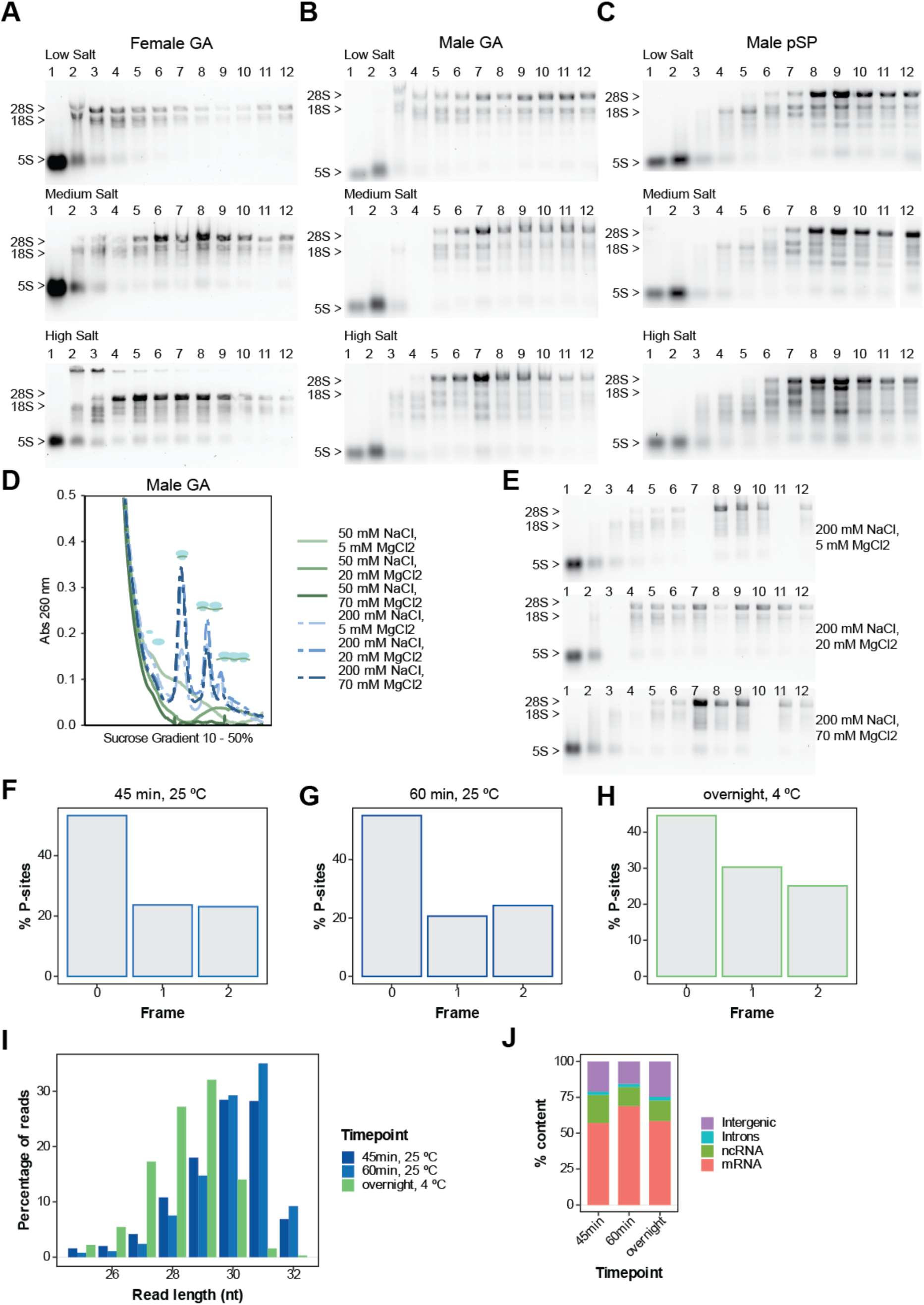
Optimization of lysis and nuclease digestion conditions for *Ectocarpus* ribosome profiling. (A–C) RNA integrity of sucrose gradient fractions assessed by electrophoresis on 1.2% agarose gels for **(A)** female GA, **(B)** male GA, and **(C)** male pSP under low (50 mM NaCl, 5 mM MgCl₂), medium (200 mM NaCl, 20 mM MgCl₂), and high (700 mM NaCl, 70 mM MgCl₂) salt conditions. Positions of 28S, 18S, and 5S rRNA detected with Midori Xtra staining are indicated. **(D)** UV absorbance profiles at 260 nm of mGA following lysis with buffers containing varying NaCl and MgCl₂. Absorbance peaks representing free 40S and 60S subunits, 80S monosomes and polysomes are indicated. **(E)** Representative gel electrophoresis images depicting rRNA integrity in the gradient fractions collected in D. **(F–H)** Reading frame distribution of P-sites for male GA Ribo-seq libraries prepared using 1 U µL⁻¹ RNase I with different digestion regimes: **(F)** 45 min at 25 °C, **(G)** 60 min at 25 °C, and **(H)** overnight (18 h, 4°C) digestion. **(I)** Ribosome protected fragment (RPF) length distribution across digestion regimes. **(J)** Distribution of RPFs within different genomic features in the Ribo-seq datasets from each digestion regime.

**Supplementary Figure 2.**
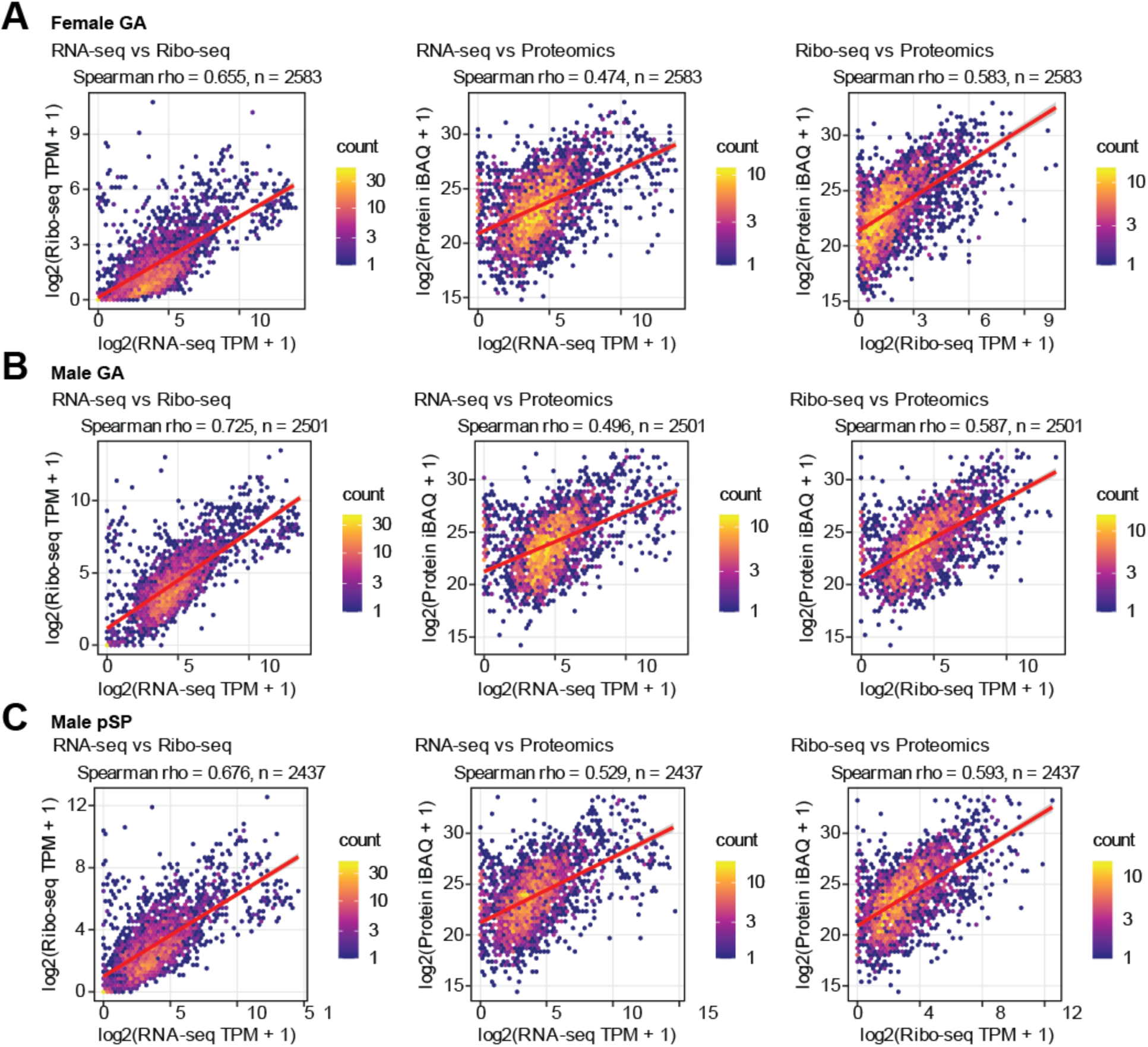
Per-stage multi-omics correlations. (A–C) Hexbin density plots showing pairwise Spearman correlations for genes detected across all three platforms in **(A)** female GA (n = 2,583), **(B)** male GA (n = 2,501), and **(C)** male pSP (n = 2,437). Left column: RNA-seq vs Ribo-seq; middle column: RNA-seq vs proteomics; right column: Ribo-seq vs proteomics. Hexagon color indicates point density. Red lines show linear regression fits. Values shown are Spearman ρ with sample sizes.

**Supplementary Figure 3.**
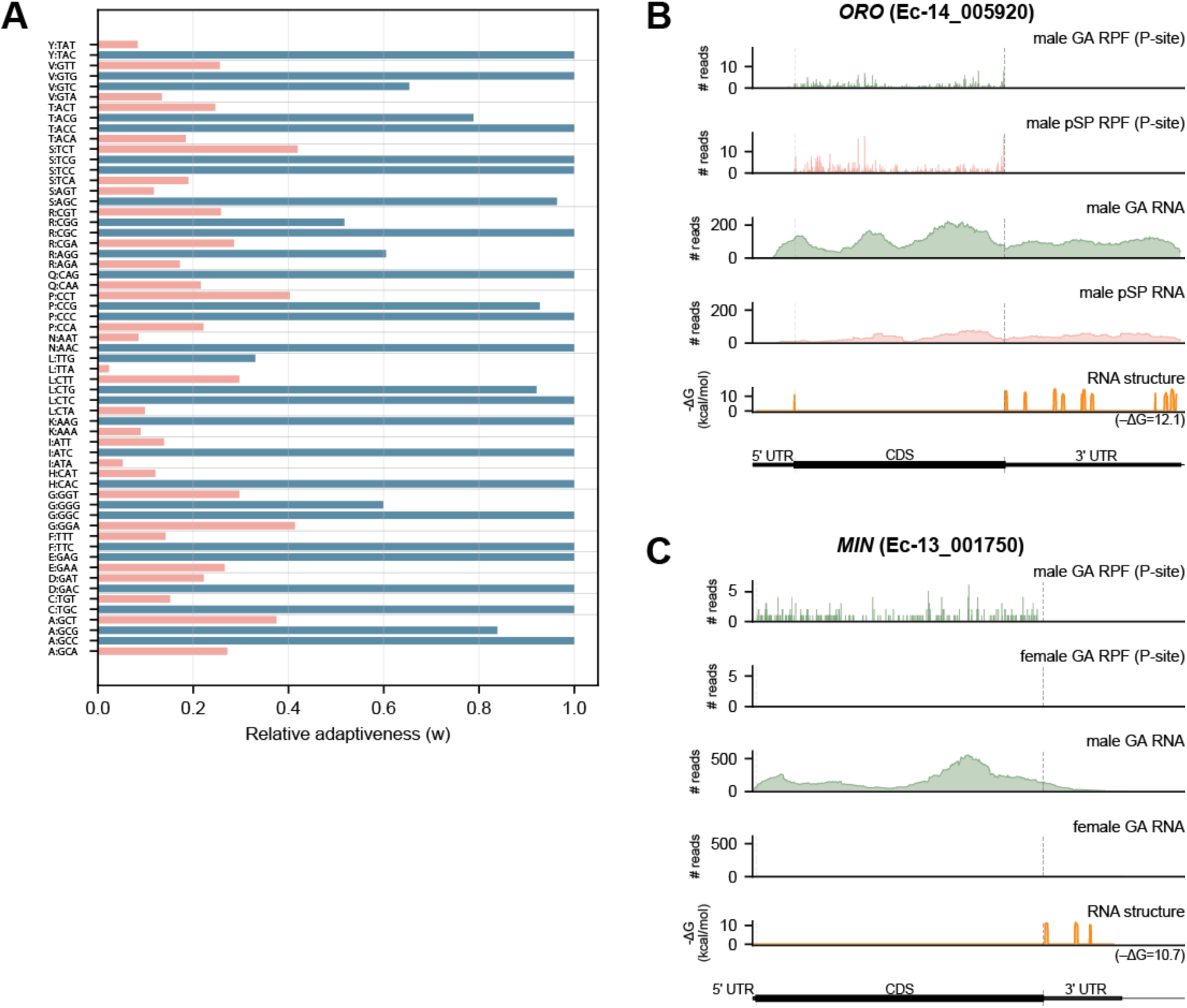
Codon usage and translational profiles of developmental regulators. **(A)** Relative adaptiveness (w) of each codon, calculated from the codon usage of the top 10% most highly translated genes (ranked by mean Ribo-seq normalized counts) using the Sharp and Li (1987) method. Codons are grouped by amino acid (one-letter code) and sorted alphabetically. For each synonymous family, w = 1 indicates the optimal codon. All 18 optimal codons are GC-ending, consistent with the GC-rich *Ectocarpus* genome. **(B)** Translational profile of *OUROBOROS* (*ORO*, Ec-14_005920), a TALE homeodomain transcription factor and master regulator of the gametophyte-to-sporophyte transition. Tracks from top to bottom: P-site-shifted Ribo-seq coverage, RNA-seq coverage, predicted RNA secondary structure propensity (–ΔG, kcal/mol, computed in sliding 40-nt windows, average value printed under the track), and gene model showing 5′ UTR, CDS, and 3′ UTR for male GA and parthenosporophyte (pSP). **(C)** Translational profile of the HMG-box sex-determination gene *MALE INDUCER* (*MIN*, Ec-13_001750). Tracks as in B, showing male and female GA comparisons. Panels B and C are plotted on the same x-axis scale to enable direct comparison of transcript length and UTR architecture between *ORO* and *MIN*.

## References

1. Hershey, J.W.B., Sonenberg, N. and Mathews, M.B. (2012) Principles of translational control: an overview. Cold Spring Harb. Perspect. Biol., 4, a011528.

2. Jackson, R.J., Hellen, C.U.T. and Pestova, T.V. (2010) The mechanism of eukaryotic translation initiation and principles of its regulation. Nat. Rev. Mol. Cell Biol. 2010 112, 11, 113–127.

3. Merrick, W.C. and Pavitt, G.D. (2018) Protein Synthesis Initiation in Eukaryotic Cells. Cold Spring Harb. Perspect. Biol., 10, a033092.

4. Brito Querido, J., Díaz-López, I. and Ramakrishnan, V. (2024) The molecular basis of translation initiation and its regulation in eukaryotes. Nat. Rev. Mol. Cell Biol., 25, 168–186.

5. Choi, J., Grosely, R., Prabhakar, A., Lapointe, C.P., Wang, J. and Puglisi, J.D. (2018) How Messenger RNA and Nascent Chain Sequences Regulate Translation Elongation. Annu. Rev. Biochem., 87, 421–449.

6. Wu, H.-Y.L., Jen, J. and Hsu, P.Y. (2024) What, where, and how: Regulation of translation and the translational landscape in plants. Plant Cell, 36, 1540–1564.

7. Liu, Y., Beyer, A. and Aebersold, R. (2016) On the Dependency of Cellular Protein Levels on mRNA Abundance. Cell, 165, 535–550.

8. Vogel, C. and Marcotte, E.M. (2012) Insights into the regulation of protein abundance from proteomic and transcriptomic analyses. Nat. Rev. Genet. 2012 134, 13, 227–232.

9. Genuth, N.R. and Barna, M. (2018) The Discovery of Ribosome Heterogeneity and Its Implications for Gene Regulation and Organismal Life. Mol. Cell, 71, 364–374.

10. Shi, Z., Fujii, K., Kovary, K.M., Genuth, N.R., Röst, H.L., Teruel, M.N. and Barna, M. (2017) Heterogeneous Ribosomes Preferentially Translate Distinct Subpools of mRNAs Genome-wide. Mol. Cell, 67, 71–83.e7.

11. Lalanne, J.B., Taggart, J.C., Guo, M.S., Herzel, L., Schieler, A. and Li, G.W. (2018) Evolutionary Convergence of Pathway-Specific Enzyme Expression Stoichiometry. Cell, 173, 749–761.e38.

12. Lorent, J., Kusnadi, E.P., van Hoef, V., Rebello, R.J., Leibovitch, M., Ristau, J., Chen, S., Lawrence, M.G., Szkop, K.J., Samreen, B., et al. (2019) Translational offsetting as a mode of estrogen receptor α-dependent regulation of gene expression. EMBO J., 38.

13. McManus, C.J., May, G.E., Spealman, P. and Shteyman, A. (2014) Ribosome profiling reveals post-transcriptional buffering of divergent gene expression in yeast. Genome Res., 24, 422–430.

14. Lyons, J., Merchante, C., Stepanova, A.N. and Alonso, J.M. (2026) Translational control in plants: from basic mechanisms to environmental and developmental responses. Plant J., 125, e70647.

15. Simsek, D. and Barna, M. (2017) An emerging role for the ribosome as a nexus for post-translational modifications. Curr. Opin. Cell Biol., 45, 92–101.

16. Saba, J.A., Liakath-Ali, K., Green, R. and Watt, F.M. (2021) Translational control of stem cell function. Nat. Rev. Mol. Cell Biol., 22, 671–690.

17. Teixeira, F.K. and Lehmann, R. (2019) Translational Control during Developmental Transitions. Cold Spring Harb. Perspect. Biol., 11, a032987.

18. Kronja, I., Yuan, B., Eichhorn, S.W., Dzeyk, K., Krijgsveld, J., Bartel, D.P. and Orr-Weaver, T.L. (2014) Widespread Changes in the Posttranscriptional Landscape at the *Drosophila* Oocyte-to-Embryo Transition. Cell Rep., 7, 1495–1508.

19. 19. Bai, B., Schiffthaler, B., van der Horst, S., Willems, L., Vergara, A., Karlström, J., Mähler, N., Delhomme, N., Bentsink, L. and Hanson, J. (2022) SeedTransNet: a directional translational network revealing regulatory patterns during seed maturation and germination. J. Exp. Bot., 74, 2416–2432.

20. 20. Bringloe, T.T., Starko, S., Wade, R.M., Vieira, C., Kawai, H., De Clerck, O., Cock, J.M., Coelho, S.M., Destombe, C., Valero, M., et al. (2020) Phylogeny and Evolution of the Brown Algae. Crit. Rev. Plant Sci., 39, 281–321.

21. Denoeud, F., Godfroy, O., Cruaud, C., Heesch, S., Nehr, Z., Tadrent, N., Couloux, A., Brillet-Guéguen, L., Delage, L., Mckeown, D., et al. (2024) Evolutionary genomics of the emergence of brown algae as key components of coastal ecosystems. Cell, 187, 6943–6965.e39.

22. Coelho, S.M., Peters, A.F., Müller, D. and Cock, J.M. (2020) Ectocarpus: An evo-devo model for the brown algae. EvoDevo, 11, 1–9.

23. Batista, R.A., Wang, L., Bogaert, K.A. and Coelho, S.M. (2024) Insights into the molecular bases of multicellular development from brown algae. Development, 151, dev203004.

24. Cock, J.M., Godfroy, O., Macaisne, N., Peters, A.F. and Coelho, S.M. (2014) Evolution and regulation of complex life cycles: a brown algal perspective. Curr. Opin. Plant Biol., 17, 1–6.

25. Coelho, S.M. (2024) The brown seaweed Ectocarpus. Nat. Methods, 21, 363–364.

26. Lipinska, A., Cormier, A., Luthringer, R., Peters, A.F., Corre, E., Gachon, C.M.M., Cock, J.M. and Coelho, S.M. (2015) Sexual Dimorphism and the Evolution of Sex-Biased Gene Expression in the Brown Alga Ectocarpus. Mol. Biol. Evol., 32, 1581–1597.

27. Cossard, G.G., Godfroy, O., Nehr, Z., Cruaud, C., Cock, J.M., Lipinska, A.P. and Coelho, S.M. (2022) Selection drives convergent gene expression changes during transitions to co-sexuality in haploid sexual systems. *Nat*. Ecol. Evol., 6, 579–589.

28. Barrera-Redondo, J., Lipinska, A.P., Liu, P., Dinatale, E., Cossard, G., Bogaert, K., Hoshino, M., Craig, R.J., Avia, K., Leiria, G., et al. (2025) Origin and evolutionary trajectories of brown algal sex chromosomes. *Nat*. Ecol. Evol., 10.1038/s41559-025-02838-w.

29. Ingolia, N.T., Ghaemmaghami, S., Newman, J.R.S. and Weissman, J.S. (2009) Genome-wide analysis in vivo of translation with nucleotide resolution using ribosome profiling. Science, 324, 218–223.

30. McGlincy, N.J. and Ingolia, N.T. (2017) Transcriptome-wide measurement of translation by ribosome profiling. Methods, 126, 112–129.

31. delCardayré, S.B. and Raines, R.T. (1995) The extent to which ribonucleases cleave ribonucleic acid. Anal. Biochem., 225, 176–178.

32. Gerashchenko, M.V. and Gladyshev, V.N. (2017) Ribonuclease selection for ribosome profiling. Nucleic Acids Res., 45, e6.

33. Meindl, A., Romberger, M., Lehmann, G., Eichner, N., Kleemann, L., Wu, J., Danner, J., Boesl, M., Mesitov, M., Meister, G., et al. (2023) A rapid protocol for ribosome profiling of low input samples. Nucleic Acids Res., 51, e68–e68.

34. Lauria, F., Tebaldi, T., Bernabò, P., Groen, E.J.N., Gillingwater, T.H. and Viero, G. (2018) riboWaltz: Optimization of ribosome P-site positioning in ribosome profiling data. PLOS Comput. Biol., 14, e1006169.

35. Tjeldnes, H., Labun, K., Torres Cleuren, Y., Chyżyńska, K., Świrski, M. and Valen, E. (2021) ORFik: a comprehensive R toolkit for the analysis of translation. BMC Bioinformatics, 22, 1–16.

36. Chothani, S., Adami, E., Ouyang, J.F., Viswanathan, S., Hubner, N., Cook, S.A., Schafer, S. and Rackham, O.J.L. (2019) deltaTE: Detection of Translationally Regulated Genes by Integrative Analysis of Ribo-seq and RNA-seq Data. Curr. Protoc. Mol. Biol., 129, e108.

37. Kryuchkova-Mostacci, N. and Robinson-Rechavi, M. (2017) A benchmark of gene expression tissue-specificity metrics. Brief. Bioinform., 18, 205–214.

38. Yanai, I., Benjamin, H., Shmoish, M., Chalifa-Caspi, V., Shklar, M., Ophir, R., Bar-Even, A., Horn-Saban, S., Safran, M., Domany, E., et al. (2005) Genome-wide midrange transcription profiles reveal expression level relationships in human tissue specification. Bioinformatics, 21, 650–659.

39. Lotharukpong, J.S., Zheng, M., Luthringer, R., Liesner, D., Drost, H.-G. and Coelho, S.M. (2024) A transcriptomic hourglass in brown algae. Nature, 635, 129–135.

40. Wright, F. (1990) The ‘effective number of codons’ used in a gene. Gene, 87, 23–29.

41. Sharp, P.M. and Li, W.H. (1987) The codon Adaptation Index--a measure of directional synonymous codon usage bias, and its potential applications. Nucleic Acids Res., 15, 1281–1295.

42. Arun, A., Coelho, S.M., Peters, A.F., Bourdareau, S., Pérès, L., Scornet, D., Strittmatter, M., Lipinska, A.P., Yao, H., Godfroy, O., et al. (2019) Convergent recruitment of TALE homeodomain life cycle regulators to direct sporophyte development in land plants and brown algae. eLife, 8.

43. Coelho, S.M., Godfroy, O., Arun, A., Corguillé, G.L., Peters, A.F. and Cock, J.M. (2011) OUROBOROS is a master regulator of the gametophyte to sporophyte life cycle transition in the brown alga Ectocarpus. Proc. Natl. Acad. Sci. U. S. A., 108, 11518–11523.

44. Luthringer, R., Raphalen, M., Guerra, C., Colin, S., Martinho, C., Zheng, M., Hoshino, M., Badis, Y., Lipinska, A.P., Haas, F.B., et al. (2024) Repeated co-option of HMG-box genes for sex determination in brown algae and animals. Science, 383.

45. Brar, G.A., Yassour, M., Friedman, N., Regev, A., Ingolia, N.T. and Weissman, J.S. (2012) High-resolution view of the yeast meiotic program revealed by ribosome profiling. Science, 335, 552–557.

46. Wu, H.-Y.L., Ai, Q., Teixeira, R.T., Nguyen, P.H.T., Song, G., Montes, C., Elmore, J.M., Walley, J.W. and Hsu, P.Y. (2024) Improved super-resolution ribosome profiling reveals prevalent translation of upstream ORFs and small ORFs in Arabidopsis. Plant Cell, 36, 510–539.

47. Blevins, W.R., Tavella, T., Moro, S.G., Blasco-Moreno, B., Closa-Mosquera, A., Díez, J., Carey, L.B. and Albà, M.M. (2019) Extensive post-transcriptional buffering of gene expression in the response to severe oxidative stress in baker’s yeast. Sci. Rep. 2019 91, 9, 1–11.

48. Ganoza, M.C., Kiel, M.C. and Aoki, H. (2002) Evolutionary Conservation of Reactions in Translation. Microbiol. Mol. Biol. Rev., 66, 460–485.

49. Mohammad, F., Green, R. and Buskirk, A.R. (2019) A systematically-revised ribosome profiling method for bacteria reveals pauses at single-codon resolution. eLife, 8, e42591.

50. Bourdareau, S., Tirichine, L., Lombard, B., Loew, D., Scornet, D., Wu, Y., Coelho, S.M. and Cock, J.M. (2021) Histone modifications during the life cycle of the brown alga Ectocarpus. Genome Biol., 22, 12.

51. Ratchinski, P., Godfroy, O., Noel, B., Aury, J.-M. and Cock, J.M. (2025) Life-cycle-related gene expression patterns in the brown algae. eLife, 14.

52. Luthringer, R., Cormier, A., Ahmed, S., Peters, A.F., Cock, J.M. and Coelho, S.M. (2014) Sexual dimorphism in the brown algae. Perspect. Phycol., 1, 11–25.

53. Kusnadi, E.P., Timpone, C., Topisirovic, I., Larsson, O. and Furic, L. (2022) Regulation of gene expression via translational buffering. Biochim. Biophys. Acta BBA - Mol. Cell Res., 1869, 119140.

54. Teyssonniere, E.M., Shichino, Y., Mito, M., Friedrich, A., Iwasaki, S. and Schacherer, J. (2024) Translation variation across genetic backgrounds reveals a post-transcriptional buffering signature in yeast. Nucleic Acids Res., 52, 2434–2445.

55. 55. Rao, S., Le, A.Y., Persyn, L. and Cenik, C. (2026) Translational buffering tunes gene expression in mice and humans. 10.1101/2025.05.16.654561.

56. Manning, J., Garcia, M.U., iraiosub, CEastwood, Tierney, J., Krueger, F., bot, nf-core, andreirie and Patel, H. (2025) nf-core/riboseq: nf-core/riboseq v1.2.0 - Mercury Spider. 10.5281/zenodo.17804124.

57. Patel, H., Manning, J., Ewels, P., Garcia, M.U., Peltzer, A., Hammarén, R., Botvinnik, O., Talbot, A., Hanssen, F., Sturm, G., et al. (2025) nf-core/rnaseq: nf-core/rnaseq v3.22.2 - Perfect Palladium Penguin. 10.5281/zenodo.17909656.

58. 58. Krueger, F. (2025) FelixKrueger/TrimGalore.

59. 59. CGATOxford/UMI-tools (2026).

60. Kopylova, E., Noé, L. and Touzet, H. (2012) SortMeRNA: fast and accurate filtering of ribosomal RNAs in metatranscriptomic data. Bioinformatics, 28, 3211–3217.

61. Liu, P., Vigneau, J., Craig, R.J., Barrera-Redondo, J., Avdievich, E., Martinho, C., Borg, M., Haas, F.B., Liu, C. and Coelho, S.M. (2024) 3D chromatin maps of a brown alga reveal U/V sex chromosome spatial organization. Nat. Commun. 2024 151, 15, 9590-.

62. Dobin, A., Davis, C.A., Schlesinger, F., Drenkow, J., Zaleski, C., Jha, S., Batut, P., Chaisson, M. and Gingeras, T.R. (2013) STAR: ultrafast universal RNA-seq aligner. Bioinformatics, 29, 15–21.

63. Patro, R., Duggal, G., Love, M.I., Irizarry, R.A. and Kingsford, C. (2017) Salmon provides fast and bias-aware quantification of transcript expression. Nat. Methods, 14, 417–419.

64. Ritter, A., Goulitquer, S., Salaün, J.P., Tonon, T., Correa, J.A. and Potin, P. (2008) Copper stress induces biosynthesis of octadecanoid and eicosanoid oxygenated derivatives in the brown algal kelp Laminaria digitata. New Phytol., 180, 809–821.

65. Cox, J. and Mann, M. (2008) MaxQuant enables high peptide identification rates, individualized p.p.b.-range mass accuracies and proteome-wide protein quantification. Nat. Biotechnol., 26, 1367–1372.

66. Tyanova, S., Temu, T. and Cox, J. (2016) The MaxQuant computational platform for mass spectrometry-based shotgun proteomics. Nat. Protoc., 11, 2301–2319.

67. Alexa, A. and Rahnenführer, J. topGO: Enrichment Analysis for Gene Ontology. doi:10.18129/B9.bioc.topGO.

68. Jones, P., Binns, D., Chang, H.-Y., Fraser, M., Li, W., McAnulla, C., McWilliam, H., Maslen, J., Mitchell, A., Nuka, G., et al. (2014) InterProScan 5: genome-scale protein function classification. Bioinformatics, 30, 1236–1240.

69. Hofacker, I.L., Fontana, W., Stadler, P.F., Bonhoeffer, L.S., Tacker, M. and Schuster, P. (1994) Fast folding and comparison of RNA secondary structures. Monatshefte Für Chem. Chem. Mon., 125, 167–188.

70. 70. Wickham, H. (2016) ggplot2 Springer International Publishing, Cham.

